# Capsid Restructuring Activates Semi-Conservative dsRNA Transcription in Cystovirus ɸ6

**DOI:** 10.1101/2025.07.23.666269

**Authors:** Serban L. Ilca, Xiaoyu Sun, Esa-Pekka Kumpula, Katri Eskelin, David I. Stuart, Minna M. Poranen, Juha T. Huiskonen

**Affiliations:** New York Structural Biology Center, New York, NY, USA; Simons Electron Microscopy Center, New York, NY, USA; Division of Structural Biology, Wellcome Centre for Human Genetics, University of Oxford, Oxford, UK; Molecular and Integrative Biosciences Research Programme, Faculty of Biological and Environmental Sciences, University of Helsinki, Helsinki, Finland; Institute of Biotechnology, Helsinki Institute of Life Science HiLIFE, University of Helsinki, Helsinki, Finland

**Keywords:** cystovirus, phi6, semi-conservative transcription, cryo-EM, cryogenic electron microscopy, localized reconstruction, symmetry relaxation, virus, dsRNA virus, bacteriophage, initiation of transcription

## Abstract

Double-stranded (ds)RNA viruses replicate and transcribe their genome within a proteinaceous viral capsid to evade host cell defenses. While *Reovirale*s members use conservative transcription, most dsRNA viruses, including cystoviruses, utilize semi-conservative transcription, where the positive strand of the genome functions as mRNA. Here, we visualize semi-conservative transcription activation in cystovirus ɸ6 double-layered particles using cryogenic electron microscopy. We observe nucleotide-triggered disassembly of the domain-swapped outer capsid layer, subsequent expansion of the inner capsid layer, and stepwise assembly of transcription complexes at the opposing poles of the spooled dsRNA genome. These complexes consist of the viral polymerases embedded into a triskelion formed by the minor protein P7, which we show as essential for continuous transcription. The packaging hexamers proximal to the transcription sites channel the viral mRNA exit. Our results define the complex molecular pathway from the quiescent state to activated semi-conservative transcription.

## Introduction

Double-stranded (ds)RNA viruses are widespread in nature and infect a broad range of hosts spanning from bacteria to humans (Neri et al., 2022; Poranen and Bamford, 2012). Unlike in other viruses, the genome replication cycle of dsRNA viruses typically occurs within a proteinaceous capsid that permanently shields the viral genome from host factors that could mediate dsRNA-induced antiviral responses (Mertens, 2004). Within the confines of the capsid, the viral RNA-dependent RNA polymerase (RdRp) replicates and transcribes the viral genome which can comprise up to twelve segments. Eukaryotic dsRNA viruses of the *Reovirales* order (hereafter reovirads), for example rotavirus (Spencer and Arias, 1981), utilize conservative transcription, whereby the newly synthesized positive-strand RNA acts as mRNA. These viruses have a four-tunneled RdRp structure, which allows separation of the newly synthetized positive-strand from the parental negative-strand (McDonald et al., 2009). In contrast to reovirads, most other dsRNA viruses, such as cysto-, partiti-picobirna-, curvula-, and birnaviruses, utilize a semi-conservative transcription mechanism (Buck, 1978; Collier et al., 2016; Dobos, 1995; Levanova et al., 2021; Usala et al., 1980). In this case, the newly synthesized positive-strand RNA replaces the parental positive-strand, which in turn is released to the capsid exterior to function as mRNA. These viruses have a three-tunneled polymerase structure, like the RdRps of positive-sense single-stranded (+ss)RNA viruses (Venkataraman et al., 2018). Regardless of transcription mechanism, the capsids enclosing the dsRNA genome provide scaffolding for genome replication and transcription during the intracellular phase of the viral lifecycle. They generally have a similar *T*=1 architecture in which the asymmetric unit is a dimer of the major capsid protein (Abrescia et al., 2012; Luque et al., 2018). The capsid encloses several copies of the RdRp subunit and, in some species, additional viral protein factors. Among the members of the order *Reovirales* and family *Cystoviridae* this single-layered particle (SLP) is typically enclosed by a second protein shell arranged on a *T*=13 icosahedral lattice to form the double-layered particle (DLP).

Among the viruses utilizing the semi-conservative transcription mechanism, cystovirus ɸ6 infecting plant pathogenic *Pseudomonas* bacteria is best characterized. In ɸ6 virions, the genome-containing expanded SLP is enclosed by an icosahedral *T*=13 layer made of protein P8 (to form the DLP) and a lipid-protein envelope (Fig 1A)(Etten et al., 1976; Huiskonen et al., 2006; Jäälinoja et al., 2007; Vidaver et al., 1973). The genome of cystoviruses comprises three dsRNA segments, S, M, and L, which are enclosed in an icosahedral *T*=1 layer (SLP) composed of 60 asymmetric dimers of the major capsid protein P1. Hexamers of packaging NTPase P4 form turret-like structures on the P1 layer at the icosahedral five-fold vertices (de Haas et al., 1999). These hexameric helicases mediate the packaging of the (+)ssRNA genomic precursors s, m, and l into the empty SLP for intra-capsid genome replication catalyzed by the enclosed P2 RdRps (El Omari et al., 2013; Ewen and Revel, 1990; Juuti and Bamford, 1995; Kainov et al., 2003; Pirttimaa et al., 2002). ɸ6 RdRps utilize de novo initiation mechanisms and share the conserved right-hand shape structure with other viral RdRps (Makeyev and Bamford, 2000b; Monttinen et al., 2021). In the empty SLP, the RdRp stabilizes the empty compact conformation of the particles by making contacts with two neighboring five-fold vertices which are oriented inward prior to genome encapsidation (Ilca et al., 2015; Sun et al., 2018). The other internal capsid protein, minor protein P7, functions as an assembly cofactor during the formation of the empty SLP (Poranen et al., 2001) and is needed for efficient genome packaging (Juuti and Bamford, 1995; Poranen et al., 2008a). Based on earlier low-resolution cryo-EM analyses, P7 could occupy a position proximal to the RdRp in the empty SLP (Nemecek et al., 2012).

**Figure 1.**
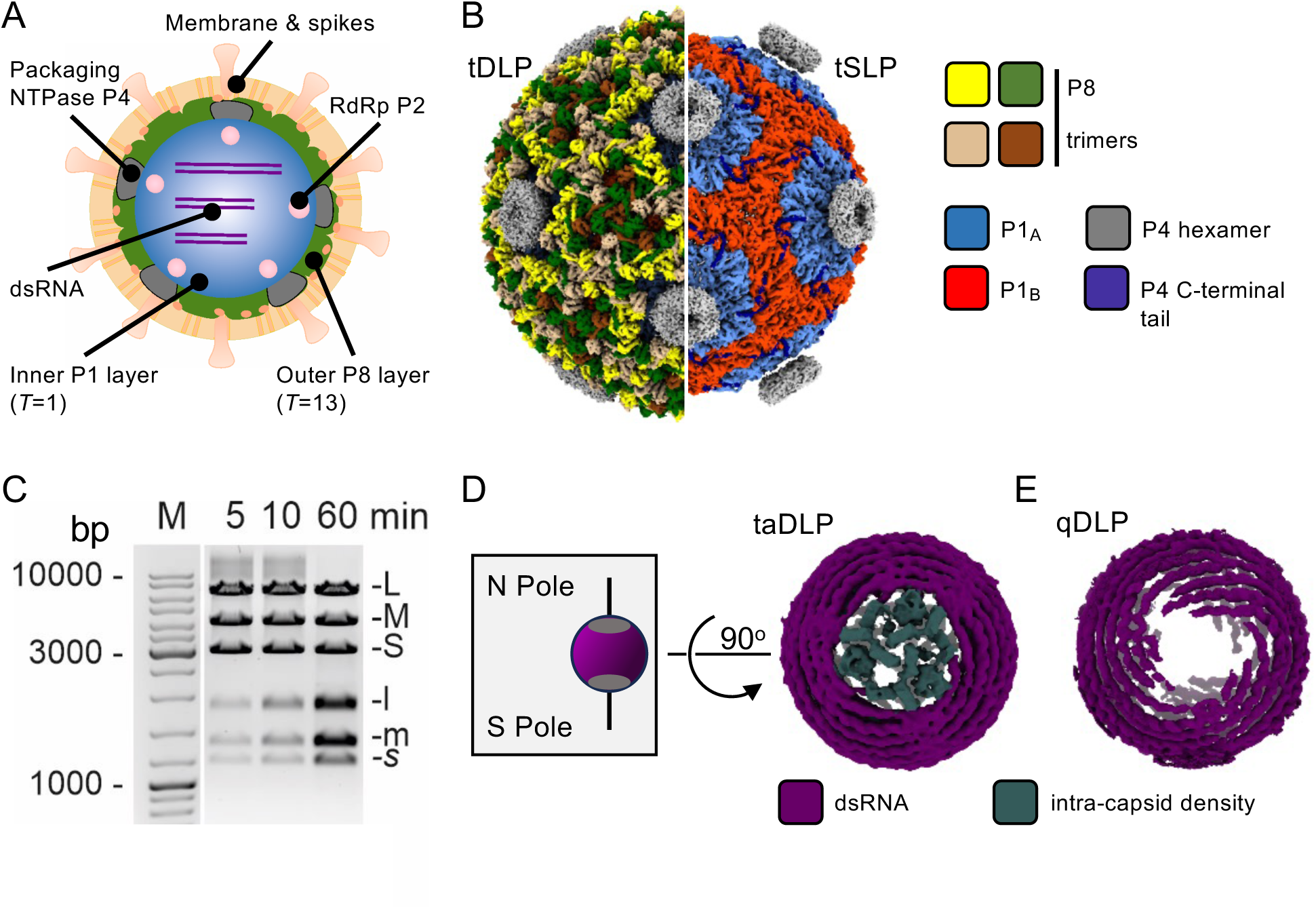
Structural reorganizations in transcribing double-layered particles. (**A**) A schematic representation of ɸ6 virion. (**B**) Cryo-EM reconstructions determined from transcribing double-layered particles (tDLPs; left) and from transcribing double-layered particles (tSLPs; right) imposing icosahedral symmetry are shown along the two-fold axis. Only half of the full cryo-EM map is shown. (**C**) Transcription activity of DLPs. Samples were taken from transcription reactions for agarose gel electrophoresis analysis at the indicated time points. The mobility of the double-stranded (uppercase letters) and single-stranded (lowercase letters) RNAs is indicated. The sizes of the molecular size marker (M) bands are indicated in base pairs (bp). (**D**) Cryo-EM reconstruction determined from transcription-arrested double-layered particles (taDLPs). The view of the genome is along the pseudo three-fold symmetry axis. Additional intra-capsid density is present at both poles of the genome (N pole and S pole), depicted in the inset (gray box). (**E**) An equivalent reconstruction as shown in (D) calculated from quiescent double-layered particles (qDLPs) published earlier (Ilca et al., 2019). The additional polar density is absent. See also Supplemental Figures 1 and 2.

In the mature DLPs, the RdRp and P7 are detached from the P1 shell, and the genome is organized as a single spool with five concentric layers around an icosahedral three-fold symmetry axis (Ilca et al., 2019). This deviates from the internal organization of the reovirads DLPs and SLPs in which the RdRp subunits are stably anchored beneath the icosahedral fivefold vertices of the inner capsid and the genome has a non-spooled configuration (Bao et al., 2022; Cui et al., 2019; Ding et al., 2019; He et al., 2019; Kaelber et al., 2020; Pan et al., 2021; Wang et al., 2018; Zhang et al., 2023; Zhang et al., 2015). These differences between reovirads and cystovirus ɸ6 suggest that the conservative and semi-conservative transcription processes of dsRNA viruses may require different intra-capsid organization. Notably, despite being widely used in dsRNA viruses, the mechanism of semi-conservative transcription and its activation remain an open question.

Here, we use ɸ6 as a model system to study the activation of semi-conservative transcription within a viral particle. Single-particle cryogenic electron microscopy (cryo-EM) reconstructions of transcribing DLPs show that the dsRNA transcription machinery, which is disordered and thus undetected in the quiescent state structure (Ilca et al., 2019), becomes ordered at the two opposing poles of the genome spool upon transcription initiation. This triskelion-shaped machinery is organized by a complex network of P7 dimers to correctly position the P2 RdRps. Using an *in vitro* assembly system to produce SLPs with reduced amounts of P7 and P2 RdRp we demonstrate that P7, in addition to P2, is essential for continuous transcription. Furthermore, the P2 RdRps contributing to genome replication are insufficient to support transcription, with additional P2s being required to assemble the P2–P7 transcription initiation complexes. The stepwise assembly of these complexes is concomitant with the NTP-triggered shedding of the outer P8 layer and further expansion of the inner P1 layer. This restructuring of the viral capsid layers is shown to be a key step in the transcription activation process and is hypothesized to play a role in the initial melting of the dsRNA template. We conclude by deriving a model for RNA template entry into the RdRp and mRNA exit from the capsid through the hexamers of P4 packaging NTPases.

## Results

### Structural reorganization in transcribing double-layered particles of cystovirus ɸ6

To study the mechanism of semi-conservative RNA transcription, we adopted a transcription-active model system based on ɸ6 doubled layered particles (DLPs)(Pirttimaa and Bamford, 2000; Van Etten et al., 1980). DLPs were prepared from purified virions by detergent solubilization of the viral membrane (see Method details). Transcription was induced by providing the four ribonucleotides (ATP, UTP, CTP, GTP), and a low-resolution structure of these transcribing DLPs (tDLP) was determined by cryo-EM (Fig 1B, Supplemental Figure 1, Supplemental Table 1). This consensus structure, calculated by averaging all particles, shows the canonical double-layered organization with P8 trimers forming the icosahedrally symmetric outer layer interrupted by 12 copies of P4 packaging motor hexamers at the five-fold vertices and P1 asymmetric dimers forming the inner layer. Closer examination revealed, however, that the outer P8 layer was incomplete or missing in ∼80% of the particles (Supplemental Figure 1A,B). Particles lacking the P8 layer were used to determine a low-resolution structure of the transcribing SLP (tSLP; Fig 1B). Under the conditions optimized for this study (see Method details), these particles efficiently transcribed all three genome segments (S, M, L) (Fig 1C), mimicking the early transcription mode of ɸ6 (Coplin et al., 1975). Full-length mRNA products (s, m, l) were present in the reaction mixture already after 5 min.

To synchronize particles in the early stages of transcription, we omitted CTP in the transcription reaction. As the first guanine in all three template strands is in the tenth position (Gottlieb et al., 1988; McGraw et al., 1986; Mindich et al., 1988), in the absence of CTP the particles are arrested after producing the first nine bases of (+)ssRNA. Analyzing a dataset of these transcription-arrested DLPs (taDLPs; 221,143 particles) allowed a thorough mapping of structural changes arising upon transcription initiation (Supplemental Figure 2; Supplemental Table 1). Similar to tDLPs, many taDLP particles showed incomplete P8 layers (Supplemental Figure 1C,D). The density corresponding to the dsRNA genome underneath the P1 shell follows the same spooled organization with pseudo-D3 symmetry as reported before for the quiescent DLP (qDLP) structure (Fig 1D,E)(Ilca et al., 2019). However, the areas at the two poles of the outmost genome layer are occupied by density in the taDLPs, whereas in qDLPs these areas are vacant (Fig 1D,E) (Ilca et al., 2019). This suggests that in transcription permissive conditions, the remaining structural components, namely the P2 RdRp and the minor capsid protein P7, become ordered. The limited resolution of these components in the cryo-EM reconstruction calculated from the complete taDLP data set indicates that different structural states were averaged together.

### Nucleotides trigger P8 outer layer shedding

To differentiate possible structural states in transcription-arrested particles, we turned to the classification of cryo-EM data. We combined 3D classification with symmetry-relaxation (Supplemental Figure 2), which allowed us first to discern the structural deviations from the icosahedral symmetry in the two protein layers. This allowed the determination of multiple cryo-EM maps showing variable degrees of P8 layer density indicating partial or complete shedding of the P8 layer in some particles (Fig. 2A–E). The shedding of P8 trimers starts with the trimers surrounding the P4 hexamers (Fig 2B,C). These P8 trimers are less tightly bound, as they interact with only two of the neighboring P8 trimers via domain-swapping, whereas trimers in the other positions swap domains with three neighbors (Sun et al., 2017). In some cases, we also detected the detachment of single P4 hexamers in the areas of P8 shedding (Fig 2B,D). In a subset of particles (∼44%), the P8 layer was fully or nearly fully absent. This subset was further used to calculate an icosahedral reconstruction of transcription-arrested single-layered particles (taSLPs; Fig. 2F; Supplemental Table 1).

**Figure 2.**
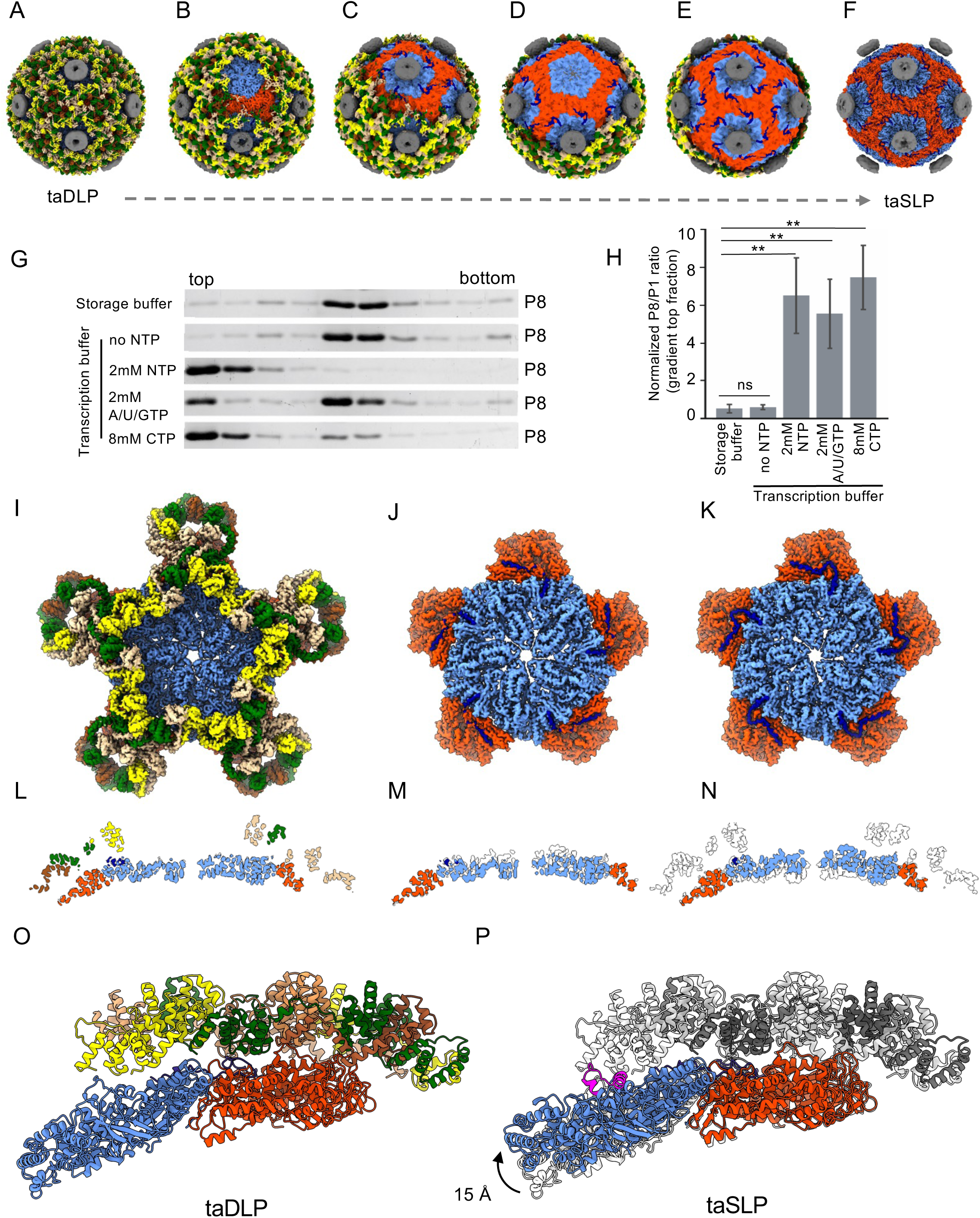
Sequential shedding of the outer P8 layer and over-expansion of the inner P1 layer. (**A**) A cryo-EM reconstruction determined from transcription-arrested double-layered particles (taDLPs). Icosahedral symmetry has been imposed on the reconstruction which is shown along the two-fold axis of symmetry. (**B–E**) Three-dimensional cryo-EM class averages of particles without symmetry, showing decreasing (dashed line) coverage of P8. (**F**) An equivalent reconstruction as shown in (A), determined from transcription-arrested single-layered particles (taSLPs) that have shed the P8 layer. The coloring of the reconstructions is as in Fig 1B. (**G**) Sedimentation analysis of the P8 layer shedding. DLPs incubated in storage buffer (no NTPs) or transcription buffer with the indicated NTPs for 10 min (the time point for taDLP cryo-EM sample preparation) were analyzed by rate-zonal centrifugation and SDS-polyacryl amide gel electrophoresis of the fractions. (**H**) The relative amount of P8 released calculated from the quantitated P1 and P8 intensities in the top fraction and plotted as P8/P1 ratio. The P8/P1 ratio observed in ɸ6 virion (analyzed in the same gel) was used for normalization. Error bars represent standard deviations of the means for four replicates. Statistically significant differences (p<0.01) between the control (no NTP) and each treatment are indicated (**). (**I,J,K**) Localized reconstructions of the double-layered shell (I), inner layer (J) and over-expanded inner layer (K) are shown. Five-fold symmetry has been imposed on all three maps to visualize the vertex assembly. The P4 hexamer density is not shown for clarity. (**L,M,N**) A slab of density is shown for the same maps as in (I), (J) and (K), rotated 90 degrees. The white overlay in (M), shown for comparison, corresponds to the map in (N). The white overlay in (N) corresponds to the map in (L). (**O**) The atomic model of the P1 asymmetric dimer fitted in the transcription-arrested double-layered particle (taDLP) cryo-EM reconstruction, together with the P8 chains proximal to it, is rendered from the side as ribbons. (**P**) The same fitting of P1 as in (O) is shown for the transcription-arrested single-layered (taSLP) cryo-EM reconstruction. The coloring of the chains is as in Fig 1B. The position of the P1 and P8 chains in the taDLP is shown as a white ribbon in the background. The upward movement of P1_A_ chain in the taSLP is indicated with an arc. The clashes this movement would cause with P8 chains are rendered in magenta. See also Supplemental Figures 2 and 3.

What is the trigger for the sequential shedding of the P8 layer? To address this, we analyzed the amounts of free P8 by rate-zonal centrifugation after incubation of DLPs in different conditions (Fig. 2G). In the qDLP controls (incubated in storage or transcription buffer), free P8 is mainly absent in the gradient top fraction (Fig. 2G; Supplemental Figure 3) showing that little P8 shedding occurs in these quiescent particles and that the P8 layer is stable in the absence of ribonucleotides. In the presence of all four ribonucleotides (ATP, UTP, CTP and GTP), 90% of the P8 layer sheds off in 10 min (Fig. 2G,H). In the transcription-arrested sample without CTP, a comparable amount of P8 shedding is observed in conditions used for cryo-EM (Fig. 2G,H). Notably, P8 shedding was also evident when the particles were incubated with CTP only (Fig. 2G,H). As these particles cannot initiate transcription, we conclude that P8 shedding is triggered by the presence of NTPs in the transcription buffer, even in the absence of transcription activation. This is consistent with earlier results showing that adding 1 mM GTP to ɸ6 DLPs increased the exposure of one P1 epitope and several P4 epitopes on the particle surface (Ojala et al., 1994).

### Outer P8 layer shedding leads to the over-expansion of the inner P1 layer

Next, we compared the state of the P1 layer in taDLPs and taSLPs. The P1 layer is known to expand during packaging of viral ssRNA into the preformed empty SLP and subsequent intra-capsid replication (negative-strand synthesis) of ssRNA to form dsRNA; this leads to the expanded mature state of the P1 shell observed in genome-containing SLPs and DLPs derived from virions (Butcher et al., 1997; Huiskonen et al., 2006). In most of the taSLPs, the P1 layer is in an over-expanded state characterized by the further bulging out of the P1 monomers adjacent to the icosahedral five-folds (type A monomers, P1_A_; Supplemental Movie 1). Little movement was observed in the other P1 monomers (type B monomers, P1_B_) relative to the taDLPs or qDLPs.

We speculated that the overexpanded state of the P1 layer may be sterically incompatible with the P8 layer assembly. Two possible but mutually exclusive scenarios could explain this: i. NTPs induce P1 layer expansion, which subsequently triggers P8 shedding, or ii. NTPs act directly on the P8 layer, inducing its shedding and allowing the P1 layer to expand spontaneously. To distinguish these possibilities, we determined the cryo-EM structures of particle surface patches for taDLPs and taSLPs by localized reconstruction (Supplemental Figure 2; Supplemental Table 2)(Abrishami et al., 2021; Ilca et al., 2015). While only the mature expanded state was observed in taDLPs (Fig. 2I,L), taSLPs exhibited both the mature and over-expanded states (Fig. 2J,K,M,N). The cryo-EM map of the P1 dimer in an over-expanded state was determined at 3.5-Å resolution, allowing us to model the P1 conformation in this state. The tip of the P1_A_ monomer, which is proximal to the five-fold axis of symmetry, swings up by 15 Å (Fig. 2P). This raised position of P1_A_ is incompatible with the P8 layer assembly (Fig. 2O), as it would cause extended clashes (Fig. 2P). As we observe uncoated patches of the P1 layer that have not yet been over-expanded (Fig. 2M), we conclude that the P8 layer acts as a harness constraining the P1 layer and that the NTP-triggered P8 layer shedding is a prerequisite for the P1 shell over-expansion.

### Minor protein P7 forms a triskelion around the P2 RdRps in transcribing particles

To understand the structural changes leading to transcription initiation in more detail, we turned our focus back to the polar regions of taDLPs and taSLPs where additional density, absent in qDLPs (Ilca et al., 2019), was observed under the P1 layer in the cryo-EM structure (Fig. 1D). To overcome the possible incoherent averaging of distinct structural states caused by the pseudo-D3 symmetry of this structure, we applied the localized reconstruction method. This method enabled the analysis of sub-particles centered at the two opposing poles from all particles. Symmetry relaxation combined with classification (Abrishami et al., 2021; Huiskonen, 2018; Ilca et al., 2019) allowed the separation of different structural arrangements of components in these polar regions (Supplemental Figure. 2). Assignment of components based on published structures of ɸ6 P2 RdRps (Butcher et al., 2001) and cystovirus ɸ12 P7 dimer (Eryilmaz et al., 2008) revealed up to three P2 RdRps per pole, in addition to a triskelion-like network of P7 dimers surrounding the P2s (Fig. 3A,B). A full complement includes four asymmetric P7 dimers per one P2 RdRp at each pole of taSLPs, totaling 24 P7 monomers and 3 P2 RdRps at each pole, and 48 P7 monomers and 6 P2 RdRps in each particle (Fig. 3B). To further support the direct P2–P7 interaction observed in these cryo-EM structures, we tested if P2:P7 complex formation could be detected in solution using purified proteins and asymmetric flow field flow fractionation (AF4). The proteins eluted together, confirming complex formation. The measured molar mass indicated that one P2 monomer and one P7 dimer form a stable 1:1 complex (Fig. 3C). Together, these results show that the organization of the internal minor capsid proteins P2 and P7 changes upon transcription and suggest an active role for P7 in the process.

**Figure 3.**
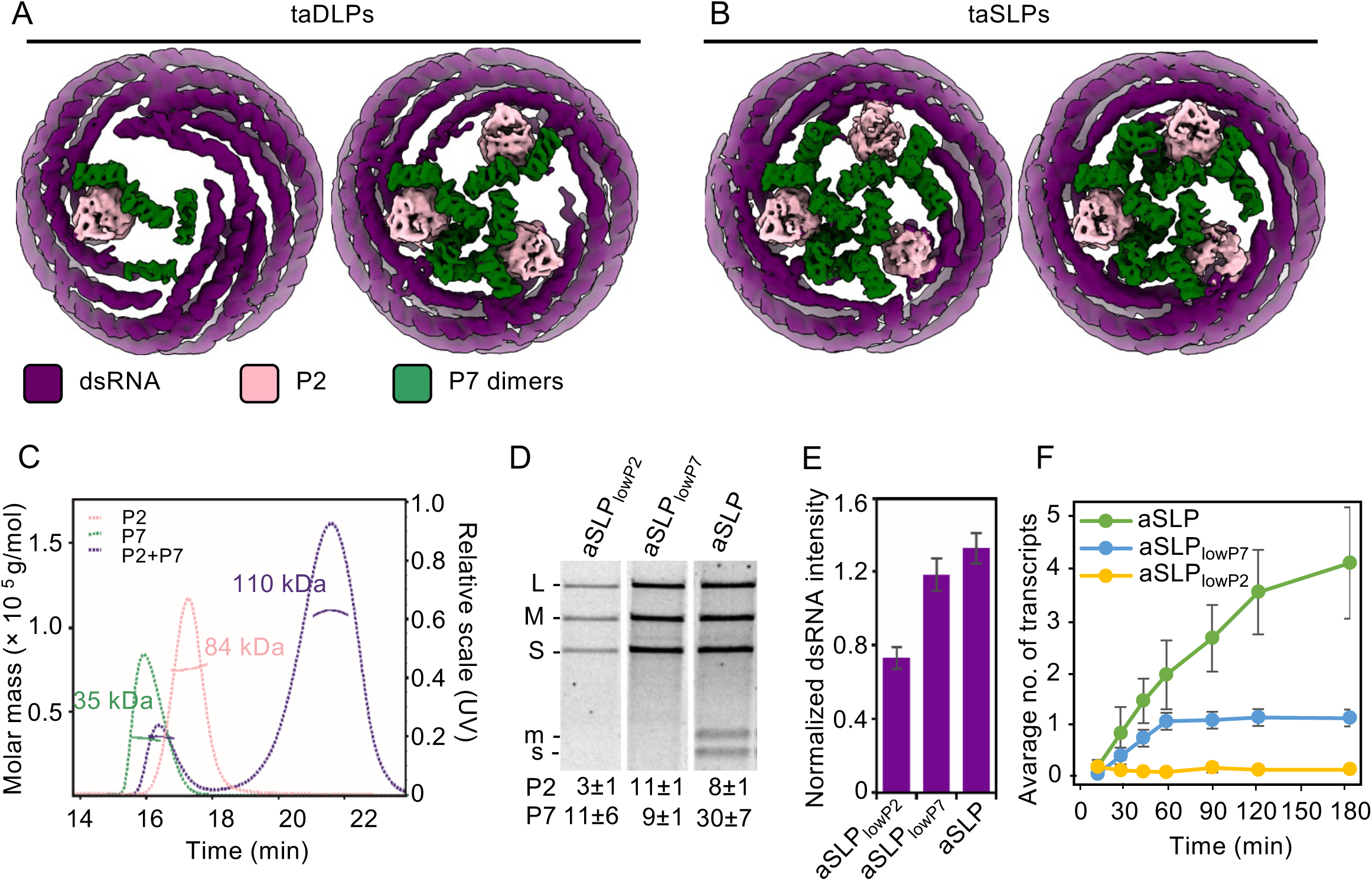
Role of P7 in the triskelion assembly and transcription. (**A**) Two three-dimensional cryo-EM class averages are shown for the polar regions extracted from transcription-arrested double-layered particles (taDLPs). (**B**) Same as (A), but the class averages have been determined from transcription-arrested single-layered particles (taSLPs). The cryo-EM averages have been filtered to local resolution. (**C**) Asymmetrical flow field flow fractionation of P2, P7 and their 1:1 mixture. Molar mass (g/mol; solid lines) and relative UV signal at 280 nm (dotted line) are shown. (**D**) Agarose gel electrophoresis analysis of *in vitro* ssRNA packaging, replication and transcription assay after a 90-min incubation with standard self-assembled SLP (aSLP) and SLPs with low amount of incorporated P7 (aSLP_lowP7_) or P2 (aSLP_lowP2_). The mobility of the synthesized ^33^P-labelled dsRNA (S, M, and L) and ssRNA transcripts (s and m) are indicated on the left of the autoradiogram, and the mean copy numbers of P2 and P7 in the SLPs with standard deviations are below. (**E**) Quantitation of dsRNA band intensities from the autoradiogram in (D). (**F**) Quantitation of time-dependent accumulation of ssRNA transcripts by the aSLPs. Error bars represent standard deviation of the mean. Calculation of the number of transcripts takes into account the semi-conservative nature of the process. See also Supplemental Figures 2 and 4.

### Half of the full complement of P7 is necessary and sufficient for efficient transcription

To further investigate the role of P7 in transcription, we turned to the ɸ6 in vitro self-assembly system. This system allows the production of SLP devoid of nucleic acid and with reduced amounts of P7 and P2, and subsequent activity tests with these *in vitro* assembled P7- (aSLP_lowP7_) and P2-deficient (aSLP_lowP2_) particles (Sun et al., 2012; Sun et al., 2018). This approach was selected because P7-null particles are compromised in genome packaging, which would have prevented the analysis of intra-capsid transcription (Juuti and Bamford, 1995; Poranen et al., 2008a), and we hypothesized that a small amount of P7 could potentially rescue the genome packaging activity. As a control, we produced aSLP particles using standard protein stoichiometry in the self-assembly reaction. We quantified the copy numbers of P2 and P7 in preparations of such particles. Firstly, the mean copy number of P2 was 8±1 copies in aSLPs (Supplemental Figure 4A). This is in line with the reported P2 copy number in ɸ6 virions (10±3; Sun et al., 2012) and suggests that there are additional copies of P2 to those 6 observed at the poles of transcribing DLPs. The amount of P7, on the other hand, averaging around 30 copies, was lower in these particles than what has been reported for ɸ6 virions (40±6; Sun et al., 2012) or what we observed in the DLPs (up to 48 copies; see Fig. 3B) (Supplemental Figure 4A). We note that this reduced set of P7 could assemble into one full P7 triskelion (24 copies) at one of the poles.

We analyzed the activities of aSLPs under *in vitro* conditions supporting the packaging of the three ssRNA genomic precursor molecules and subsequent intra-capsid genome replication and transcription of the small (S) and medium (M) segments (Fig. 3D,E; late transcription mode of ɸ6)(Coplin et al., 1975). The reactions were carried out in the presence of ^33^P-UTP and unlabelled viral genomic ssRNA molecules, resulting in the radioactive label incorporation first into the negative and positive strands of the dsRNAs (S, M and L) and subsequently in the ssRNA transcripts (s and m). It is worth noting that because of the semi-conservative transcription mechanisms and the use of unlabelled ssRNA, the first transcripts produced from each segment are unlabelled and, instead, the intensity of the label in the dsRNA doubles (Supplemental Figure 4B).

The aSLP control particles with 30 copies of P7 on average efficiently replicated the viral ssRNAs and supported the average production of four transcripts per replicated genome segment when the incorporation of ^33^P-UTP in nascent RNA was analyzed by agarose gel electrophoresis (Fig. 3D,E,F). The aSLP_lowP7_ and aSLP_lowP2_ particles had a reduced set of P7 (only 9 and 11 on average, respectively) but still showed genome packaging and replication activity (Fig. 3D,E). While aSLP_lowP2_ particles were transcription deficient, the dsRNA signals produced by aSLP_lowP7_ and aSLP control particles were roughly equal (Fig. 3D,E), indicating that aSLP_lowP7_ particles were able to start the first round of transcription (Fig. 3F). Interestingly, these particles produced no labeled ssRNA (Fig. 3D) indicating that these particles could not support additional rounds of transcription after the first one. This deficiency seems to be linked directly to the diminished amount of P7 in these particles, as they had a normal amount of P2 (11±1) (Fig. 3D,E; Supplemental Figure 4A). On the other hand, the SLP_lowP2_ particles, having 3±1 P2s on average, could replicate the tri-segmented viral genome, but could not transcribe at all (Fig. 3D,E,F). The lack of transcriptional activity in these particles was directly connected to the reduced amount of P2 RdRp, as these particles had similar amount of P7 as the SLP_lowP7_ particles (Fig. 3D, Supplemental Figure 4C).

In conclusion, around half of the full complement of P7 (approximately thirty copies in the aSLP control particles) is necessary and sufficient for multiple rounds of transcription, presumably by allowing the assembly of one complete transcription complex from twenty-four P7 monomers at one pole. Furthermore, particles having three P2s could replicate the tri-segmented viral genome but could not transcribe. This suggests that two pools of P2s are present. One pool (with as few as three P2s) is necessary and sufficient for replication. A second additional P2 pool is required to assemble a functional transcription complex composed of three P2 RdRps and twenty-four P7 monomers.

### Stepwise assembly of the transcription complex

How does the P7 triskelion assemble? And does the triskelion assembly order the three P2 RdRps, or do they organize independently? We addressed these questions by localized reconstruction of cryo-EM data where contributions of P1 and P8 layers had been subtracted. We centered the analysis on each individual P2 and classified the density corresponding to P2, P7 and surrounding dsRNA (Supplemental Figure 2; Supplemental Table 3). We observed different structural states, differing in the number of ordered P2 and P7 proteins and the amount of ordered RNA. These states were ordered as three stages (A, B, C), the first having three sub-stages (A1–A3), starting from least to most ordered (Fig. 4A). In the first sub-stage (A1), both P2 and an asymmetric dimer of P7 dimers are present, suggesting that they co-assemble, as neither partner is observed alone in these data. This agrees with our observation that P2:P7 complex forms in solution (see above; Fig. 3C). This stage is followed by the second dimer of P7 dimers becoming ordered in the following sub-stages (A2 and A3). This second dimer of dimers is in a conformation (“wrapped”) differing from the conformation of the first (“extended”). In stage B, additional density is present, which we assign to RNA. This density is in direct contact with the wrapped P7 dimer of dimers and becomes more prominent in the final stage C. We speculate this density may correspond to the 3’-end of the (-)ssRNA template, a notion that requires further verification. Consistent with this hypothesis, the P7 C-terminus of a related cystovirus ɸ12 shows affinity to ssRNA (Eryilmaz et al., 2008).

**Figure 4.**
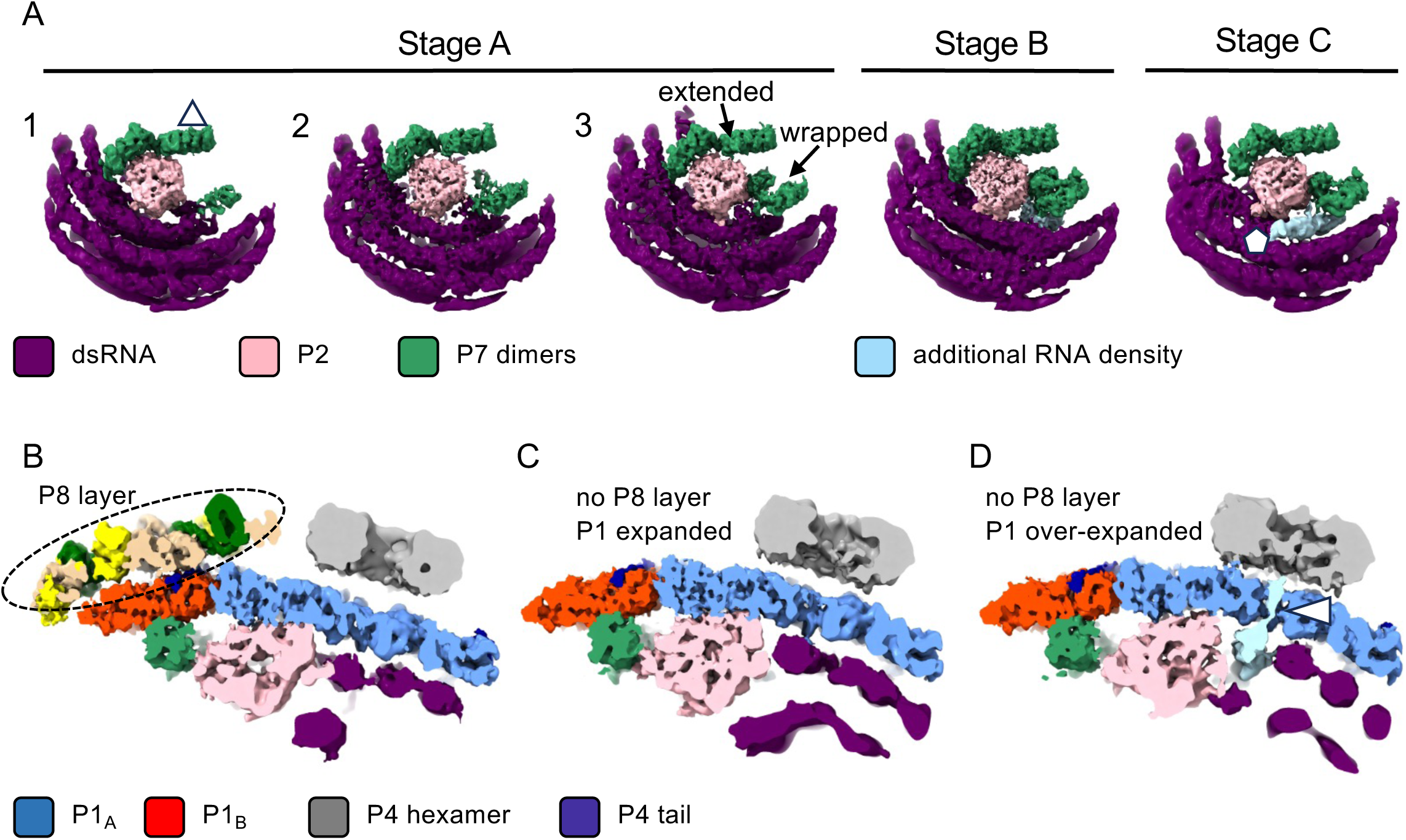
Stepwise assembly of the transcription initiation complex. (**A**) Five three-dimensional cryo-EM class averages (stages A1, A2, A3, B and C) are shown for the local P2 neighborhoods, extracted from transcription-arrested particles. The local three-fold axis of symmetry is indicated (triangle) in stage A1. Two conformations of the P7 dimer of dimers are seen in stage A3, “extended” and “wrapped”. The latter is partially disordered in stages A1 and A2. The maps have been filtered to local resolution and segmented manually to remove inner and outer protein layers for clarity. (**B**, **C**, **D**) The same maps, as shown in panel (A) stages A1, B, and C, respectively, are shown from the side. The outer P8 layer (circled with a dashed line) is present in stage A1, but absent in the two other stages. In the final stage shown in panel (D), the P1 layer is over-expanded and putative RNA density (light blue) extends through the P1-shell opening at the five-fold vertex (white arrowhead). See also Supplemental Figure 2.

To study the possible link between P2/P7 transcription complex assembly and P1 layer expansion/P8 layer shedding, we analyzed the structural state of the P1 and P8 layers in the three stages by calculating cryo-EM maps by localized reconstruction (Fig. 4B–D). In stages A1–A3, the P8 layer is still present (Fig. 4B), whereas in stages B and C, it is absent (Fig. 4C,D). These two stages differ in the conformation of the P1 shell; in stage B, the P1 shell is similar to the normal expanded state, whereas in stage C, the P1 shell is clearly overexpanded (see also Fig. 2N). In stage C, the putative dsRNA end seems split, as narrow density is observed extending through the pore at the five-fold position in the P1 layer towards the central cavity of the P4 hexamer (Fig. 4D). This density likely corresponds to the 5’-end of the (+)ssRNA, that is, the mRNA to be extruded from the particle. In conclusion, these results show that two asymmetric dimers of P7 dimers are organized around each P2 RdRp, suggesting that P7 may act as an actuator of transcription initiation. Furthermore, the final stages of P7/P2 triskelion assembly, whereby P7 interacts with the putative template RNA strand, are linked to the shedding of the outer P8 layer, the over-expanded state of the P1 layer, and positioning of the second RNA strand for particle exit.

### P7 orchestrates the arrangement of RdRps to the transcription initiation sites

To understand the detailed interactions between P7 and other components of the particle, we first modeled the atomic structure of the P7 dimer (residues 2–112 of the total 161 residues) based on the localized reconstruction of the polar region at an average resolution of 4.1 Å and an AlphaFold3 prediction of two copies of the complete P7 sequence (Fig. 5A–C)(Abramson et al., 2024). Similar to the cystovirus ɸ12 P7 (Fig. 5D)(Eryilmaz et al., 2008), the ɸ6 P7 consists of a central parallel four-stranded β-sheet (β1–β4), surrounded by three shorter helices (⍺2–⍺4) and a long C-terminal helix (⍺5). The N-terminal helix in ɸ12 P7 (⍺1), proposed to stabilize the overall fold via interactions with the C-terminal helix (Eryilmaz et al., 2008), is absent in ɸ6 P7. Instead, ɸ6 P7 contains an additional helix between β1 and β2 (denoted here as ⍺1’). Key differences were also detected in the P7 dimerization interface. Whereas in the crystal structure of ɸ12 P7, the dimerization surface involves a predominantly hydrophobic patch of nine residues from ⍺2 and ⍺3, the ɸ6 P7 dimerization interface shows a stacking interaction between non-conserved tryptophan residues (W37 in ⍺3; Fig. 5E).

**Figure 5.**
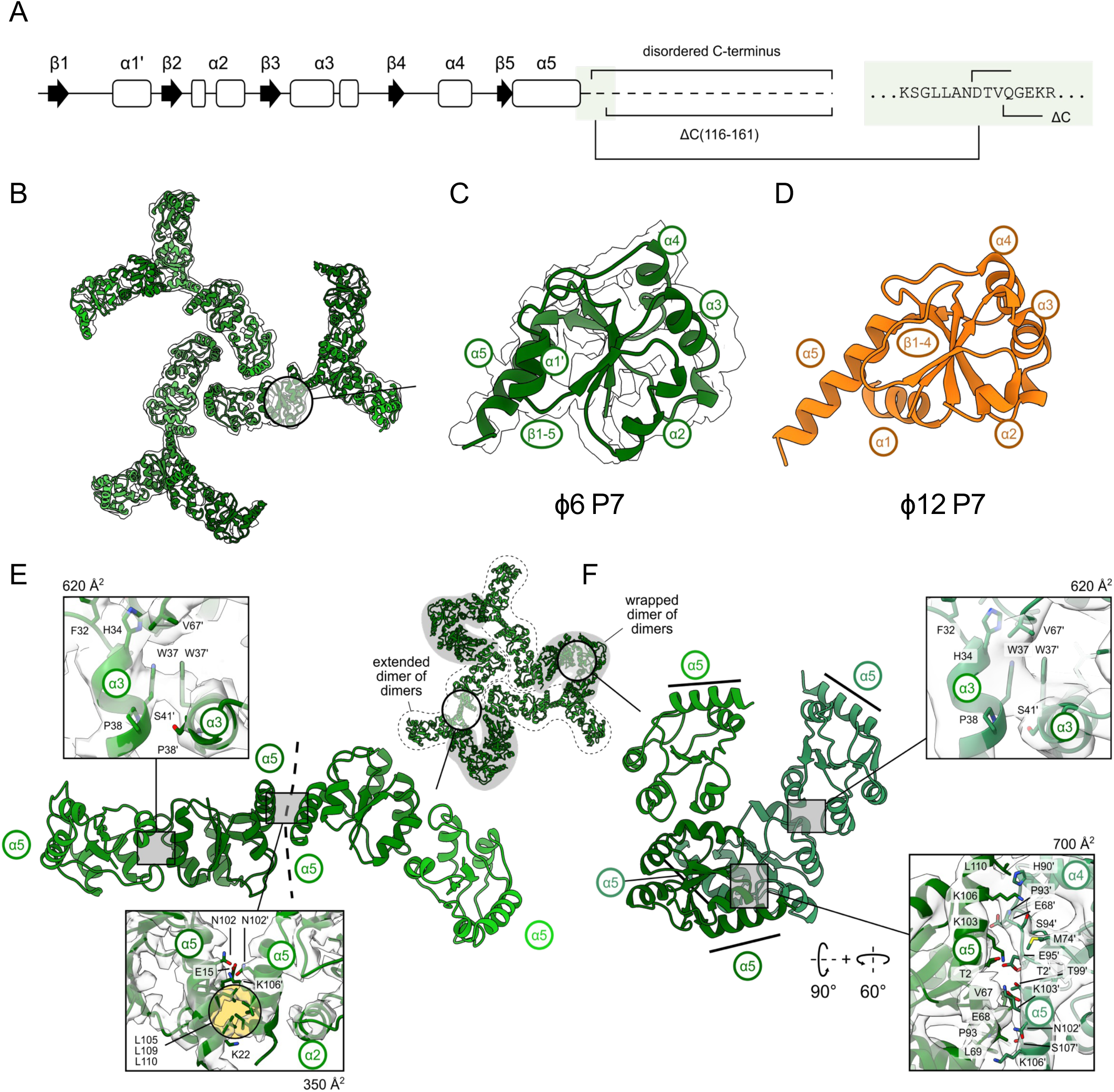
The structure of minor protein P7. (**A**) Secondary structure elements of the P7 monomer. The C-terminal truncation site is indicated in the inset. (**B**) Overview of the P7 triskelion model within cryo-EM density map low-pass filtered to 6-Å resolution. A region corresponding to approximately one monomer is circled. (**C**) Model of ɸ6 P7 monomer fitted into highest resolution region of the consensus P7 triskelion reconstruction. (**D**) Model of ɸ12 P7 monomer (PDB:2Q82). (**E**) The architecture of the P7 dimer interfaces in the extended dimer of dimers. The close-ups show the monomer–monomer (top) and the dimer–dimer (bottom) interfaces. Dashed line indicates the extended dimer-dimer interface. The inset (top, right) shows the arrangement of the extended dimers of dimers (outlined with a dashed line) and wrapped dimers of dimers (gray background). (**F**) P7 dimer interfaces in the wrapped dimer of dimers. The close-ups show the monomer–monomer (top; cryo-EM density map low-pass filtered to 6-Å resolution) and the dimer–dimer (bottom) interfaces. The approximate interaction interface surface areas are given. See also Supplemental Figures 2, 6 and 7.

To model the complete assembly, the P7 dimer model was fitted as a rigid body to all P7 positions in our highest resolution cryo-EM map (stage B) (Fig. 6A,B). The dimer–dimer interface in both the extended (Fig. 5E) and wrapped (Fig. 5F) dimer of dimers is mediated by ⍺5. However, the mode of interaction differs significantly due to the significant difference between the two conformations.

**Figure 6.**
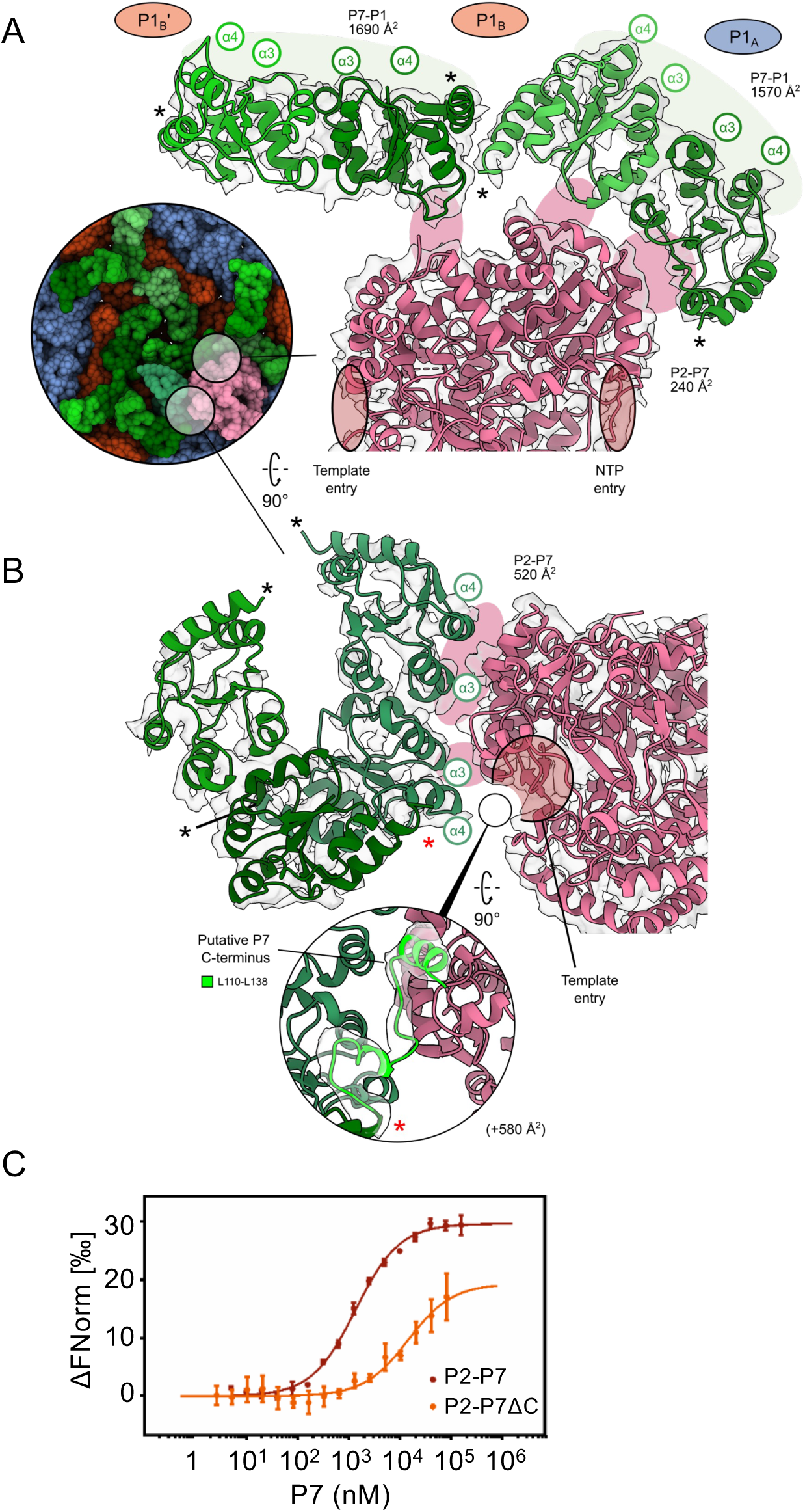
Interactions of minor protein P7 with the inner P1 shell and P2 RdRp. (**A**) The orientation of P7 extended dimer of dimers (green) relative to the inner P1 shell and the P2 RdRp (pink). (**B**) The interaction interface of P2 and wrapped P7 dimer of dimers. The close-up shows a putative interaction of the P7 C-terminus (L110–L138) proximal to the template entry site of P2. The P7 model with C-terminal residues predicted by AlphaFold3 was flexibly fit into the cryo-EM density map low-pass filtered to 6-Å resolution. The C-termini of the P7 models are denoted with asterisks. The openings of the template and NTP tunnels on the RdRp surface are indicated. Interaction interfaces in both (A) and (B) are indicated with light green or pink background for P7-P1 and P2-P7 interfaces, respectively, in addition to the corresponding interface area estimates. (**C**) Microscale thermophoresis analysis of P2 interaction with full-length P7 (red) or P7ΔC (orange). The error bars represent the standard deviation of each data point calculated from three measurements. See also Supplemental Figures 2, 6 and 7.

The fitting allowed the mapping of four extensive interaction sites between P7 and the inner surface of the P1 layer (Supplemental Figure 5; sites a1, a2, b1 and b2). The extended P7 dimers of dimers that assemble to the transcription initiation site already at stage A1 has a “basket”-like structure on one side of the dimer (formed by ⍺3 and ⍺4 helices), which interacts with P1_B_ chains around the three-fold axis of symmetry (Supplemental Figure 5C, sites b1 and b2; approximately 290 Å^2^ and 1700 Å^2^, respectively). Extended P7 dimers of dimers (monomers A–D) also show a secondary interaction surface to the P1_A_ chains around the five-fold axes of symmetry (site a1; 310 Å^2^). The wrapped dimer of dimers that is fully assembled only after stage A3, on the other hand, interacts solely with the P1_A_ chains (site a2; 690 Å^2^). In addition to the P7 basket, sequence motifs (denoted here as curvature-specific elements) mediate interactions with the P1 layer depending on its specific local curvature (Supplemental Figure 5C,D). We suggest that these interactions of the P7 dimers with P1 prime them to multimerize further and position P2 RdRps in their correct locations close to the five-fold axes of symmetry proximal to each pole.

How does P7 interact with P2 RdRp? A previously published atomic structure of P2 (Butcher et al., 2001) was fitted as a rigid body to the stage B cryo-EM map. This fitting unambiguously established the orientation of the P2 relative to the P1 layer and P7 triskelion, and some of its contacts with the ordered RNA (Fig 6; Supplemental Figure 6). Despite apparent surface complementarity and proximity in all stages A–C (Fig. 3A,B), only a few interactions were detected between the extended P7 dimer of dimers and P2 by ChimeraX interface analysis (Fig. 6A). However, the wrapped P7 dimer of dimers interacts with P2 via ⍺3 and ⍺4 (approximately 520 Å^2^), in addition to a putative C-terminal helix-loop-helix motif (Fig. 6B). Cryo-EM density for this motif, missing in the initial P7 model (covering residues 2–112; Fig. 5), was observed on the side of P2 proximal to the template entry site and modeled by AlphaFold3. This motif is predicted to contribute an additional interaction surface measuring approximately 580 Å^2^.

In solution, P7 is partially disordered (Benevides et al., 2002) and sensitive to proteolytic cleavage close to the C-terminal end (at residue 120)(Poranen et al., 2008a). We suggest that when P7 is bound to P2 in the transcriptionally active SLP, the C-terminal part becomes more ordered, forming a C-terminal helix-loop-helix motif. This is not only consistent with the cryo-EM density but also with earlier observations of increased helicity in the capsid-associated state of P7 (Benevides et al., 2002). To test this proposed interaction further in solution, we produced C-terminally truncated P7 comprising residues 2–115 (P7ΔC) and analyzed binding to P2 using microscale thermophoresis (MST). The measured dissociation constant of P7ΔC (15.1±3.4 µM) was significantly higher than for the full-length P7 (1.3±0.1 µM) (Fig. 6C). Together, these experiments confirm that the P7 C-terminal domain from the wrapped dimer of dimers contributes to P2 binding. In conclusion, via its interactions with the P1 layer, P7 orchestrates the assembly of the transcription initiation site by positioning the P2 RdRps correctly to support transcription.

### Hexameric P4 packaging motors act as passive conduits for mRNA release

How do the mRNA transcripts exit the transcribing particles? Cryo-EM reconstructions of taDLPs and taSLPs with D3 symmetry show that the central pore in the “polar” P4 hexamers, which are proximal to the transcription complexes, is open in taDLPs but filled in taSLPs (Fig. 7). This is consistent with the localized reconstruction of stage C transcription initiation complex, also showing density extending towards the P4 central pore (see Fig. 4D). What is the significance of this occluding density? Each hexamer can assume any of the five possible poses relative to the underlying five-fold symmetric protein layer beneath it. As a result, hexamers with different poses are averaged together in the reconstruction, and their density becomes obscured. To overcome this C5–C6 symmetry mismatch effect (Huiskonen, 2018), we applied symmetry-relaxation on the P4 hexamer sub-particles extracted by localized reconstruction. This allowed detailed analysis of the hexamer structure, including the occluding density. First, we focused on hexamers extracted from taDLPs (Fig. 7A). The equatorial and polar P4 hexamers bound to the P1 layer were both resolved to 4.9-Å average nominal resolution (Supplemental Table 2). In both cases, the central channel of the P4 hexamer was open (Fig. 7B–E). Next, we carried out the analysis with P4 hexamers extracted from taSLPs (Fig. 7F). The equatorial P4 hexamers display an open pore at 5.2-Å average nominal resolution (Fig. 7G,H). For polar P4 hexamers, however, a comparison of the structural states revealed that in one class at 6.0-Å average nominal resolution, the pore is occluded by a 73-Å long tubular density that cannot be explained by the presence of P1 or P4 proteins in the vicinity and which we thus assign as mRNA (Fig. 7I,J). These results are consistent with earlier studies showing that while a single P4 hexamer is sufficient to support genome packaging into empty SLPs, additional hexamers are required to support the transcription (Pirttimaa et al., 2002; Sun et al., 2013). Furthermore, while energy from NTP hydrolysis by P4 is needed for ssRNA packaging into empty SLPs, it is not needed for the nascent transcript export (Kainov et al., 2004). Taken together, these results show that the central pores of P4 hexamers act as passive conduits necessary for mRNA release from the taSLPs.

**Figure 7.**
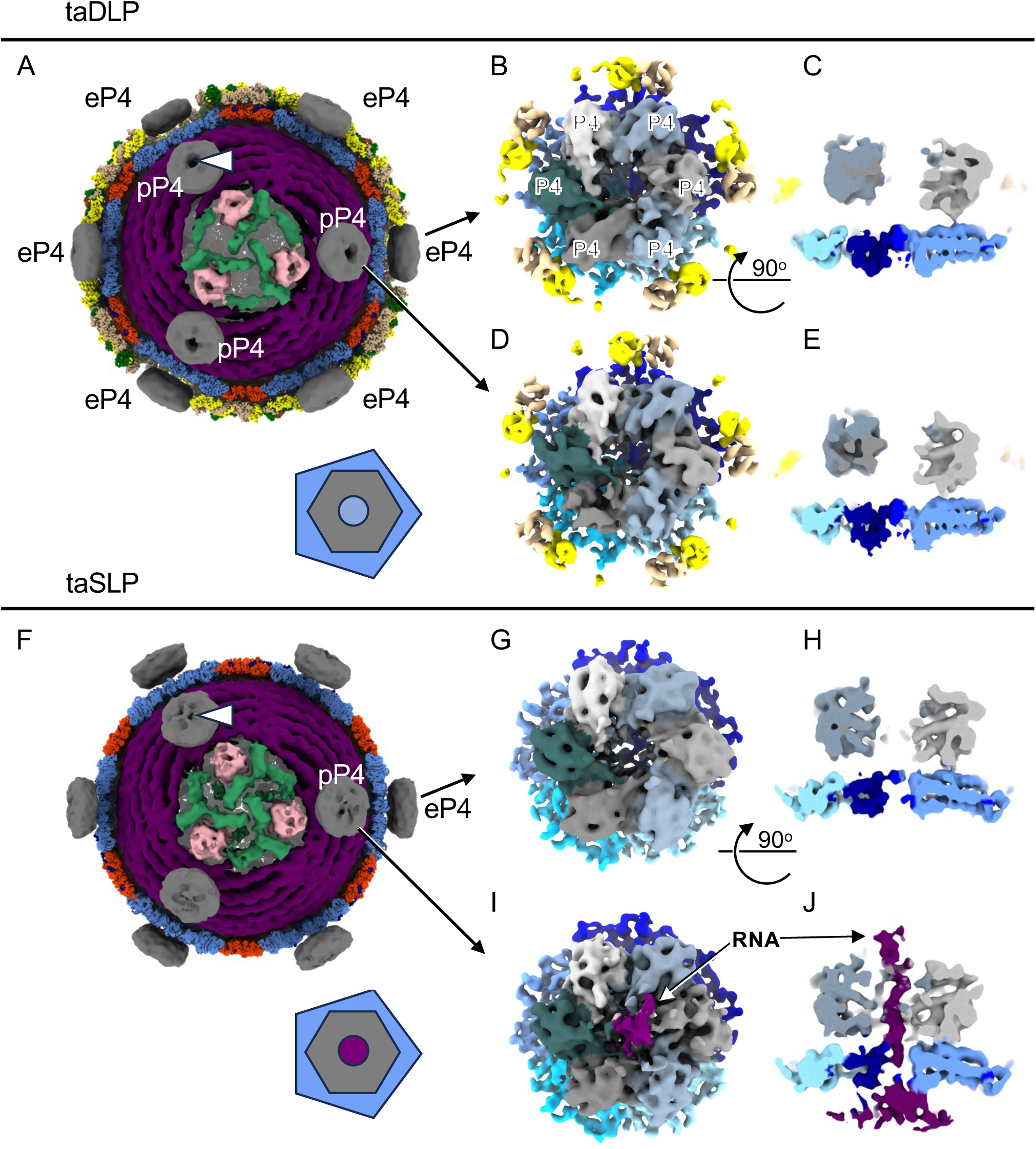
P4 hexamer acts as a conduit for mRNA release. (**A**) A cryo-EM reconstruction determined from transcription-arrested single-layered particles (taDLPs) with D3 symmetry shown along the three-fold axis of symmetry. The protein shells have been removed in the front half to show the genome and the polar region. Three P4 hexamers, adjacent to the polar region and the P2 RdRps (pink), are labeled (pP4). The central cavity in these polar P4 hexamers appears plugged (arrowhead). The six P4s, distal to the polar region (equatorial P4s), are labeled (eP4). The inset illustrates the symmetry mismatch between the hexamer (grey hexagon) and the underlying P1 shell (blue pentagon). (**B,C**) Three-dimensional cryo-EM class average of equatorial P4 hexamers from taDLPs is shown from top (B) and side (C). Individual P4 monomers are labeled in (B). (**D,E**) Class average of polar P4 hexamers from taDLPs. (**F**) The equivalent reconstruction as shown in (A) determined from transcription-arrested single-layered particles (taSLPs). The central cavity in the polar P4s appears plugged (arrowhead), also illustrated in the inset (purple circle). (**G,H**) Three-dimensional cryo-EM class average of equatorial P4 hexamers from taSLPs is shown from top (G) and side (H). (**I,J**) Class average of polar P4 hexamers from taSLPs. Additional density, assigned as RNA (purple) is seen to plug the P4 cavity. Coloring is as in Fig 1B. See also Supplemental Figure 2.

## Discussion

Most dRNA viruses perform semi-conservative intra-capsid transcription which had been poorly understood. Here we studied transcriptionally active cystovirus ɸ6, which utilizes semi-conservative transcription, and observed RdRp subunits clustered around two opposite three-fold symmetry positions at both poles of the spooled genome. These RdRps are proximal to the neighboring five-fold vertices and embedded in a complex triskelion-like structure formed by dimers of the minor capsid protein P7. The observed intra-capsid organization of the minor capsid proteins in transcribing and arrested DLPs differs significantly from their placement in quiescent DLPs and unexpanded empty SLPs (Ilca et al., 2015; Ilca et al., 2019; Nemecek et al., 2012), highlighting extensive structural changes leading to transcription initiation.

Our data allows us to propose a model for the activation of semi-conservative transcription in ɸ6 (Fig. 8). The fully assembled P8 layer sequesters SLP in a transcriptionally inactive state and transcription initiation is accompanied by the melting of the dsRNA end, enabling mRNA exit at the five-fold vertex through the hexameric P4 packaging NTPase. All previously reported triggers of P8 outer layer shedding, namely either low pH or a Ca^2+^ chelating agent (Cvirkaite-Krupovic et al., 2010; Olkkonen et al., 1991), were absent in the conditions used here. Instead, we observed that NTPs in the transcription buffer act as the trigger for outer layer shedding, even in the absence of transcription initiation. The pattern of P8 shedding suggests a domain-unswapping mechanism that starts from the P8 trimers surrounding the P4 hexamers with fewer stabilizing domain-swaps than the other P8 trimers (Sun et al., 2017). Previous time-resolved measurements on ɸ6 DLPs have identified local unfolding events, characterized by low activation barriers in the P8 layer (Tuma et al., 1999). Based on the identification of multiple P8 disassembly intermediates, we postulate that the assembly/disassembly reaction adheres to the principles of a biochemical equilibrium driven by NTP concentration. As such, if the NTP concentration in the bacterium is high, P8 disassembly is favored, resulting in continued viral replication. Conversely, if the NTP concentration is low and Ca^2+^ ions are available, P8 assembly is favored, resulting in viral maturation and exit.

**Figure 8.**
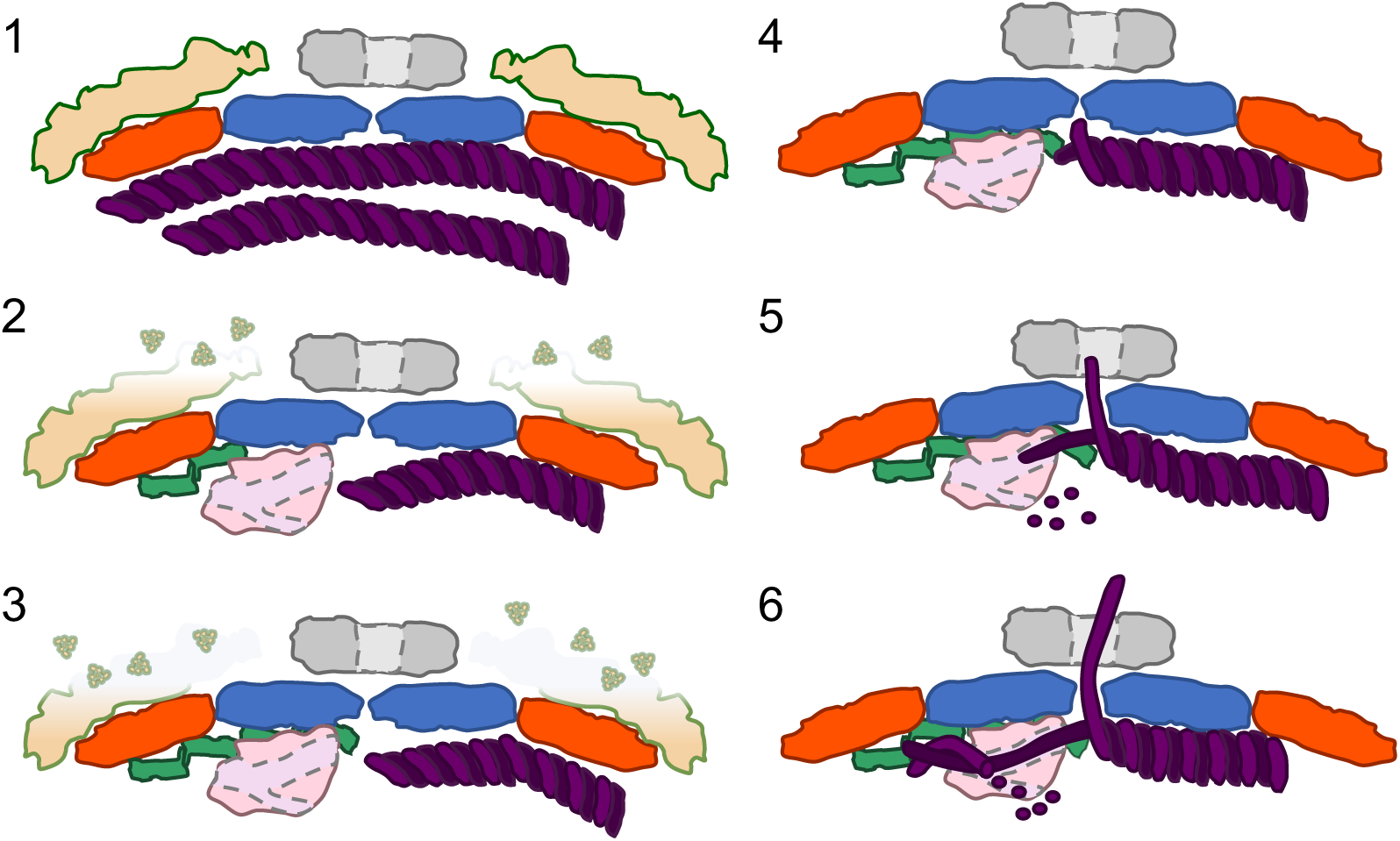
Model for activating semi-conservative transcription in ɸ6. A schematic depiction of the transcription initiation complex assembly (steps 1–6). Step 1: Quiescent state of the double-layered particle with no ordered P2 or P7. Step 2: Partial shedding of the outer P8 layer (beige), partial assembly of the P7 triskelion (green) with ordered P2 (pink). Step 3: Further shedding of the outer P8 layer and increased copy number of P7. Step 4: Fully shed P8 layer, fully assembled P7 triskelion with the “wrapped” P7 dimer, bringing the end of the dsRNA template (purple) to the polymerase. P1 layer (red and blue) is still in its normal state. Step 5: Overexpansion of the P1 layer leads to the melting of the dsRNA template. Step 6: Active transcription and the exit of the transcript through the P4 (grey) central pore.

Following P8 layer shedding we observe over-expansion of the inner P1 layer proximal to the transcription sites. The mechanism of this expansion remains unknown, and it can be driven either by the potential energy stored in the particle during assembly or by energy released by NTP hydrolysis at the early stage of transcription. As RdRps in solution are inefficient in initiating transcription (Makeyev and Bamford, 2000a), we hypothesize that the over-expansion of the P1 layer occurs spontaneously and drives the melting of the dsRNA template, providing a free 3’-end of the negative-strand template for transcription. This hypothesis is supported by a long ssRNA strand exiting the particle without any dsRNA duplex exiting from the P2 RdRp during this stage. This density (73 Å in length) would correspond to a minimum of 12 nucleotides even if the ssRNA was in a fully over-stretched confirmation (assuming 5.9 Å/nt) (Smith et al., 1996). Furthermore, we note that the length of this mRNA is longer than the 9 nucleotides that the transcription-arrested particles can synthesize before stalling. After the initial melting, the RdRp can efficiently catalyze the strand displacement (Makeyev and Bamford, 2000a). Local resolution analysis of the P2 RdRp density in the cryo-EM maps of different stages indicated that the RdRps become increasingly mobile when transcription starts (Supplemental Figure 6B). Finally, the NTP-trigger may provide an elegant switch from transcription to the assembly of progeny virions. We hypothesize that the depletion of NTPs in the host cell cytoplasm at the late stage of infection switches the particles from transcription to the assembly of progeny virions, as in the presence of calcium and the absence of NTPs, the RNA synthesis is turned off, and the P8 shell can assemble on the quiescent SLPs (Olkkonen et al., 1991; Salgado et al., 2004). Testing this hypothesis would require measuring NTP and calcium levels in the host cell at different time points during infection.

Based on our results, minor protein P7 emerges as a major orchestrator of intra-capsid transcription. This adds to its previously reported roles in assembling empty ɸ6 SLPs (Poranen et al., 2001), and genome packaging (Juuti and Bamford, 1995; Poranen et al., 2008a). The ⍺-helical basket allows the P7 dimer to bind to different sites on the inner surface of the P1 shell and also the P2 RdRp (Supplemental Figure 7). Our data demonstrate that genome packaging is rescued, resulting in intra-capsid genome replication if the SLPs contain approximately 9 P7s (Fig. 3D). Such particles can also support one round of transcription from the newly synthesized dsRNA genome, while approximately 30 P7 subunits are sufficient to support continuous intra-capsid transcription. This corresponds to about half of a full complement of P7 proteins or one complete P7 triskelion assembly (24 proteins), highlighting the system’s inherent evolutionary robustness. However, despite this built-in resilience, maintaining balanced stoichiometry remains crucial. In ɸ6, this balance is achieved through regulated protein expression, facilitated by the translation of SLP components from a single polycistronic mRNA, while the relative production of P2 and P7 is further controlled by coupling their synthesis through translational reinitiation (Mindich et al., 1988). These results extend our understanding of multifunctional protein complexes and how stochiometric fine-tuning may modulate their functional properties. Furthermore, we speculate that the role of P7 in ɸ6 intra-capsid transcription is to bring the 3’-end of the genomic negative-strand and the P2 RdRp in close proximity to facilitate the initiation complex formation for positive-strand synthesis (Fig. 4A, Stage C). Such activity is specifically required for transcription re-initiation, as after positive-strand synthesis, the P2 RdRp is positioned at the 5’-end of the negative-strand distal to the 3’-end of the template strand at the opposite end of the dsRNA genome (Supplemental Figure 4B). However, after completion of genome replication, the P2 RdRp is positioned at the 3’-end of the nascent negative-strand (Supplemental Figure 4B, steps II and III) and no movement of P2 or the template is needed to initiate the production of the first transcript. This likely explains why the production of the first transcript by ɸ6 SLP is less dependent on P7 than the synthesis of the additional transcripts (Fig. 3D).

Cryo-EM combined with localized reconstruction facilitated here the *in situ* structural analysis of an RdRp in a transcribing viral particle. Previous structural, biochemical and biophysical analyses on the ɸ6 RdRp have provided detailed insights into the template and nucleotide entry into the catalytic site (Poranen et al., 2008b; Salgado et al., 2004) and formation of the initiation complex (Butcher et al., 2001), in addition to the transition to elongation (Wright et al., 2012) and the kinetics of elongation (Dulin et al., 2015). The cryo-EM data presented shows 3 copies of P2 RdRp located at both genome spool ends in transcribing particles suggesting that 3 to 6 RdRps could participate in the transcription of the tri-segmented ɸ6 dsRNA genome. However, previous biochemical data has shown that ɸ6 virions contain approximately 10 RdRp (Sun et al., 2012) suggesting that additional RdRps, which are more randomly oriented and thus unobserved in the cryo-EM maps, are present in the DLPs. We hypothesize that these additional RdRps could have a specialized role in the replication of the tri-segmented dsRNA during the maturation of the genomecontaining DLPs. In line with this, we have observed earlier that SLPs containing only 3 RdRps can catalyze the synthesis of the minus-strand for the three genomic single-stranded precursor molecules but are defective in transcription (Sun et al., 2018)(see also Fig. 3D). In this model, these 3 RdRps would not correspond to the RdRps which contribute to the formation of transcription complexes at either of the two spool ends and are required to rescue transcription. Our insights into the structural organization of the transcription complexes thus provide a framework for these earlier observations and suggest potential functional specialization for the RdRps.

The process of intra-capsid transcription initiation in ɸ6 resembles that in other dsRNA viruses. Similar to ɸ6 P8 shell, Ca^2+^ stabilizes the outer T=13 protein shell in rotavirus virions (Salgado et al., 2018). Further shared features among reovirads and ɸ6 are that the disassembly of the outer T=13 shell, if present, is required to produce the transcriptionally active particle (Danthi et al., 2010; Jayaram et al., 2004; Roy, 2020), the inner protein shell is expanded in the transcription preinitiation/initiation stage (He et al., 2019; Jenni et al., 2019; Stevens et al., 2023; Yang et al., 2012), and that the transcript likely exits through channels at the 5-fold vertex (Jenni et al., 2019; Lawton et al., 1997; Yang et al., 2012; Zhang et al., 2003). Whether the outer shell disassembly in reovirads is also triggered by the presence of NTPs remains an open question. A notable difference is that reovirads utilize conservative transcription mechanisms and the RdRps are stably positioned underneath the five-fold vertices both in the quiescent and in transcriptionally active stages, while semi-conservative transcription in cystovirus ɸ6 requires restructuring of the capsid interior and assembly of transcription complexes at the opposing poles of the spooled dsRNA genome. Further studies on partiti-picobirna-, and curvulaviruses are required to verify if such mechanism is common for dsRNA viruses using semi-conservative intra-capsid transcription. Furthermore, recent *in situ* cryo-electron tomography studies tracing dsRNA virus infection steps *in vivo* for bluetongue virus (Shah et al., 2023; Xia et al., 2024) are expected to inspire similar studies on cystoviruses.

## Acknowledgements

We acknowledge Electron Bio-Imaging Centre (eBIC) at the Diamond Light Source, Oxford Particle Imaging Center (OPIC) cryo-EM facility at the University of Oxford and Instruct Centre UK for access and support. The facilities and expertise of the HiLIFE Biocomplex unit and CryoEM unit at the University of Helsinki, members of Instruct-ERIC Centre Finland, FINStruct, and Biocenter Finland are gratefully acknowledged. The MST interaction analyses were performed at the Biomolecular Interaction Unit, Faculty of Biological and Environmental Sciences, University of Helsinki. We thank Xun Lu for assisting in model building. This study was supported by grants from the Research Council of Finland (331627 to M.M.P.), Jane and Aatos Erkko Foundation (to J.T.H. and M.M.P.), Sigrid Jusélius Foundation (220147, 230156, 240167 to M.M.P.), Ella and Georg Ehrnrooth Foundation, Finnish Cultural Foundation (to X.S.) and UK Medical Research Council (MR/N00065X/1 to D.I.S.). S.L.I. was supported by a Wellcome Trust four-year PhD studentship (109135/Z/15/A) and by the National Resource for Automated Molecular Microscopy, funded by NIH National Institute of General Medical Sciences (GM103310) and the Simons Foundation (SF349247). Work in the laboratory of J.T.H. was supported by Helsinki Institute of Life Science HiLIFE. The authors wish to acknowledge CSC – IT Center for Science, Finland, for generous computational resources.

## Author contributions

Conceptualization, M.M.P, J.T.H.; Formal Analysis, S.L.I., X.S., E-P.K., K.E.; Investigation, all authors; Writing – Original Draft, J.T.H., M.M.P.; Writing – Review & Editing, all authors; Visualization, S.L.I., E-P.K.; Funding Acquisition, S.L.I., J.T.H., M.M.P.; Resources, M.M.P., J.T.H.; Supervision, D.I.S., M.M.P., J.T.H.

## Methods

### Bacterial strains, viruses and plasmids

*Pseudomonas syrinage* pathovar *phaseolicola* HB10Y was used for the propagation of cystovirus ɸ6 (Vidaver et al., 1973). Recombinant ɸ6 proteins P2, P7, P7ΔC, and P4, were expressed from plasmids pEM2, pEM7, pSB1 and pJTJ7 (Makaya and Bamford, 2000b; Ojala et al., 1993; Poranen et al., 2008a; Poranen et al., 2001), respectively, in *Escherichia coli* BL21(DE3) or HMS174. P1P4 particles for P1 purification were produced in *E. coli* JM109(pLM358)(Gottlieb et al., 1990). Plasmids pLM659, pLM656 and pLM687 (Gottlieb et al., 1992; Mindich et al., 1994; Olkkonen et al., 1990) were used as templates for ɸ6-specific ssRNA production by T7 transcription (s, m, and l, respectively).

### Purification of DLPs

DLPs were obtained from purified ɸ6 virions by removal of the viral envelope using Triton X-114 extraction and subsequent chromatographic purification in a CIM DEAE-1 tube monolithic column (BIA Separations, Slovenia) (Sun et al., 2017). The freshly made particles in 20 mM K3PO_4_, pH 7.2, 1 mM MgCl_2_, 0.1 mM CaCl_2_, 150 mM NaCl (storage buffer) were used for transcription reactions.

### Production of empty ɸ6 SLPs and their *in vitro* activity analyses

**Recombinant SLPs (**rSLPs) were produced in *E. coli* JM109(pLM687)(Pirttimaa et al., 2002) and purified by Triton X-114 extraction and rate-zonal centrifugation (Gottlieb et al., 1990). *In vitro* self-assembly of SLPs was induced by mixing purified recombinant proteins P1, P2, P4 and P7 (Juuti and Bamford, 1997; Makeyev and Bamford, 2000b) in 50 mM Tris-HCl (pH 8), 6% (wt/vol) polyethylene glycol (PEG) 4000, and 1 mM ATP as described before (Poranen et al., 2001; Sun et al., 2012). The resulting aSLPs were separated from the proteins subunits by rate-zonal centrifugation. For the analysis of the combined in vitro ssRNA packaging, replication and transcription activity of aSLPs, the plus-strand synthesis assays were performed according to a protocol published earlier (van Dijk et al., 1995) in 50 mM Tris-HCl pH 8.9, 2 mM DTT, 0.1 mM EDTA, 5 mM MgCl_2_, 6% (wt/vol) PEG 4000, 80 mM NH_4_Ac, 20–44 ng/µl s, m and l ssRNAs in equal molar ratio, 1 mM NTP and [α-^33^P]-UTP (PerkinElmer) with 0.04 mg/ml SLP at 30°C for 15-180 min. For the time course analyses, 4 µl samples were taken at the indicated time points and immediately mixed with urea containing 2×U loading buffer (Pagratis and Revel, 1990). The reaction products were analyzed by electrophoresis in a native agarose gel and the autoradiographs were recorded using a phosphorimager (BAS-1500, Fujifilm).

### Transcription reactions with ɸ6 DLPs and analysis of P8 shell shedding

The transcription reactions were carried out in 50 mM Tris-HCl pH 8.0, 50 mM NH_4_Ac, 100 mM KCl, 5 mM DTT, 3 mM MnCl_2_, 1 mM MgCl_2_, 0.005–0.01mM CaCl_2_, 2 mM NTP (for tDLPs) or A-/G-/UTP (for taDLP) and 0.4 U/µl Ribolock RNasin (Thermo Fisher Scientific), using 0.15–0.5 mg/ml DLP in final volume of 40 µl. The reactions were incubated at 30°C for 5 to 60 min and reaction products analyzed by electrophoresis in native agarose gel. GeneRuler DNA Ladder Mix (Thermo Fisher Scientific) was used as a molecular size marker. P8 shell shedding was analyzed by ultracentrifugation in 10–30% sucrose gradient, 20 mM Tris-HCl, pH 8, 150mM NaCl, 1mM MgCl_2_ (Sorval AH650 rotor, 35,000 rpm, 25 min).

### Cryo-EM sample preparation, data collection and processing

A 3-µL aliquot of the transcription reaction with purified ɸ6 DLPs after a 10-min incubation was applied to a glow-discharged electron microscopy grid coated with a holey carbon foil (C-flat; Protochips). The grid was blotted and then vitrified in liquid ethane using Vitrobot (Thermo Fisher Scientific). Cryo-EM data on transcribing particles were collected on a Polara transmission electron microscope operated at cryogenic conditions and at 300 kV using a Gatan K2 Summit direct electron detector in counting mode and magnification of 100,000× resulting in a calibrated pixel size of 1.35 Å. Cryo-EM data on transcription-arrested particles were collected over two sessions on two different transmission electron microscopes (Titan Krios, Thermo Fisher Scientific) operated at cryogenic conditions and at 300 kV using direct electron detectors (Falcon 3, Thermo Fisher Scientific) in linear mode and magnification of 100,000× resulting in a nominal pixel size of either 1.39 Å (session 1) or 1.44 Å (session 2). Data collection parameters are summarized in Supplemental Table 1.

All data were processed in RELION 3.0.6 (Zivanov et al., 2018) within the Scipion 3.0 framework (de la Rosa-Trevín et al., 2016) unless stated otherwise. For both transcribing and transcription arrested particle datasets, the same procedures were applied as before for quiescent ϕ6 (Ilca et al., 2019) to perform motion correction, CTF estimation (Rohou and Grigorieff, 2015) and 2D classification. The transcribing particles were semi-automatically picked using Xmipp (de la Rosa-Trevín et al., 2013) and classified into five 2D classes. One class (46% of particles) corresponded to tSLPs, two classes (22% and 19%) to tDLPs and two classes (13%) to particles with a partial P8 layer. The tDLP and tSLP cryo-EM maps were reconstructed with icosahedral symmetry.

The particles in the two datasets of transcription-arrested particles were automatically picked in ETHAN (Kivioja et al., 2000) and separately classified into 50 2D classes in RELION. The first dataset led to one taSLP class (27%), two taDLP classes (16% and 9%) and one class with a partial P8 layer (10%), while the second gave two taSLP classes (16% and 9%), one taDLP class (23%) and one class with a partial P8 layer (8%). Particles with P8 (both complete and incomplete taDLPs) were separated from particles that lacked the P8 layer (taSLPs). For combining the data from two different imaging sessions, preliminary taDLP cryo-EM maps were calculated in RELION using icosahedral symmetry. The pixel sizes were calibrated by fitting an atomic structure (PDB:6HY0) in UCSF ChimeraX (Meng et al., 2023), resulting in calibrated pixel sizes of 1.36 Å and 1.40 Å. To combine the two datasets, the latter one was scaled from 1.36 Å to 1.40 Å using Xmipp (de la Rosa-Trevín et al., 2013). Cryo-EM maps of both taDLPs and taSLPs were reconstructed using icosahedral symmetry. To reconstruct the taDLPs without icosahedral symmetry for visualization, we subjected the particles to 3D classification using a mask defining the capsid layers. To relax icosahedral symmetry, we used a custom version of RELION (https://github.com/serbanilca/relion-3.1-relax). Here, only the 60 orientations related by icosahedral symmetry were probed and the resolution used in classification was limited to 60 Å (parameter “--strict_highres_exp”). Twelve distinct classes were further reconstructed without any symmetry applied to resolutions in the 6.3–20.5 Å range. Four best resolved maps (taDLP disassembly intermediates 1–4) were included in further analysis.

To reconstruct the capsid interior, a similar procedure as for qDLP (Ilca et al., 2019) was performed individually for taSLP and taDLP particles. First, the signal corresponding to the capsid was computationally subtracted from the original images. Second, 3D classification with symmetry relaxation and one class was performed using the qDLP map filtered to 60 Å as an initial model. The resulting map exhibited pseudo-D3 symmetry, so the map and the particles were rotated to follow D3 symmetry convention. Two distinct rounds of classification were performed. First, subparticles from the two spool ends were extracted and combined into a single dataset. 3D classification relaxing C3 symmetry was performed into 4 classes. Two best resolved maps for both taSLP and taDLPs, corresponding to accurately aligned particles, were included in further analysis. Second, six subparticles each corresponding to one P2 neighborhood were extracted and combined. 3D classification was performed without applying symmetry and without rotations or translations (“--skip_align”) into 10 classes and subsequent rounds of 3D classification were performed to obtain a total of 5 well-resolved maps between taSLP and taDLP. To include those components in the analysis that were subtracted earlier (P1 and P8 density), we assigned the alignment parameters of relevant subparticles for each localized reconstruction calculated above to subparticles extracted from the original (unsubtracted) particles.

For P4 processing, twelve subparticles were extracted separately for taDLP and taSLP: six corresponding to polar P4 regions and six corresponding to equatorial P4 regions. Only subparticles corresponding to side views (maximum deviation of ±30°) were retained for further processing. Classification was performed with C5 symmetry relaxation into 8 classes for the polar P4’s from taSLP and into 4 classes each for the other three types of P4’s. Finally, to further analyze the mode of RNA exit, we resolved the three-way symmetry mismatch between P1, P2 and P4 in the same vertex by subclassifying the subparticles showing RNA density into 5 classes without alignment.

The data set corresponding to the taSLP spool ends was used to determine the structure of P7. First, C3 symmetry expansion was performed, followed by focused classification using a mask corresponding to a single P7 dimer of dimers part of the P7 triskelion core. The full triskelion was obtained from the particles corresponding to the class with the most apparent P7 density features by reconstruction using the signal from the entire spool end.

The resolution in all cryo-EM maps was estimated from two independent half sets of particles using Fourier shell correlation following the gold-standard postprocessing and cut-off of 0.143 in RELION. The maps were sharpened by applying the inverse B-factor. Maps were further filtered to local resolution for visualization. Map coloring was based on the proximity to the fitted atomic models (see below) and manual segmentation for RNA. Data processing parameters are summarized in Supplemental Table 2.

### Atomic models

To model the different states of the capsid layers (DLP inner layer expanded, SLP inner layer expanded and SLP inner layer overexpanded), an atomic model of ɸ6 asymmetric unit (PDB:6HY0) was flexibly fitted in ISOLDE (Croll, 2018) within UCSF ChimeraX (Meng et al., 2023) and refined in Phenix (Liebschner et al., 2019) using the default parameters generated by command “*isolde write phenixRsrInput*”. P2 (PDB:1HHT) and P4 (PDB:5MUV) were fitted as rigid bodies in UCSF ChimeraX (Meng et al., 2023). The six subunits of P4 were handled as independent rigid bodies. The initial model of P7 was predicted using AlphaFold3 (Abramson et al., 2024) with two copies of the full sequence of P7. One P7 dimer was fitted into the P7 triskelion map at the highest resolution P7 site using ISOLDE and refined in Phenix. Atomic model of stage B was prepared as a composite of flexibly fitted P1 chains using ISOLDE and Phenix and rigid body fitted models of P7 dimers and P2.

### Measurement of protein interactions

AF4 experiments were performed using EclipseTM NEON (Wyatt Technology, Dernbach, Germany) field flow fractionation system (Eskelin et al., 2022) at 22°C using a long analytical channel, a 400-µm spacer (Wyatt Technology), a 10-kD regenerated cellulose membrane (Wyatt Technology) and 10 mM Tris-HCl (pH 8.0), 10 mM NaCl, 0.1 mM EDTA as the mobile phase. Channel and detector-flow velocities were 1.0 and 0.5 ml/min, respectively. A constant cross-flow velocity of 5 ml/min was utilized for separation. The weight-average molar mass (M) was obtained using ASTRA software v. 8.1.2 using a UV detector signal at 280 nm as the concentration detector and calculated absorption coefficients of 1.212 and 1.437 for P7 and P2, respectively. P2 and P7 complex formation was studied by mixing them in the used mobile phase in 1:1 molar ratio (P2 20 µg and P7 9 µg) in 20 µl final volume prior to the AF4 experiment.

MST measurements were performed at 50% LED power and 40% MST power at 25°C on a Monolith NT 115 instrument (NanoTemper Technologies GmbH, München, Germany). P2 labeled with RED fluorescent dye NT-647-NHS (Monolith Protein Labeling Kit RED-NHS) was used at 28–35 nM concentration and mixed with the full-length P7 or P7ΔC in 20mM Tris (pH8.0), 0.2 mg/ml BSA, 0.05% Tween 20, 10mM NaCl at room temperature. P7 or P7ΔC were titrated in 2-fold dilutions from the initial concentration of 65 µM or 86 µM, respectively. After a 10-min incubation, the protein mixtures were loaded into Monolith NT.115 Capillaries (NanoTemper Technologies) for measurements. The binding curve fitting was done using MO.Affinity Analysis v. 2.3 (NanoTemper Technologies).

### Quantification and statistical analysis

ImageJ was used for the densitometric analysis of protein band intensities in SDS-polyacrylamide gels stained with Coomassie brilliant blue and ^33^P-label intensity of RNA molecules in the autoradiographs of agarose gels. Relative amounts of proteins P1, P2 and P7 in the purified SLP samples were calculated based on the measured protein band intensities in SLPs and in ɸ6 virion sample, and the known stoichiometry of the proteins in phi6 virions as described (Sun et al., 2012). T-test was used to analyze statistical differences between measured protein ratios assuming equal variance and two-tailed distribution. The [α-^33^P]UTP-labeled dsRNA and ssRNA products were quantified from autoradiograms of agarose gels using Image J. In the calculation of relative transcript copy numbers the nature of semi-conservative transcription was taken into account. Briefly, the first round of transcription releases the unlabelled parental (+)ssRNA and the plus-strand in the dsRNA becomes labeled resulting in doubling of dsRNA label intensity (negative-strand is already labeled). The subsequent rounds of transcription produce labeled ssRNA molecules. Thus, the number of transcripts produced per each negative-strand was obtained by i) dividing the measured ssRNA intensity by the intensity of the negative strand, which was obtained by dividing the dsRNA intensity by two, and then ii) adding the value corresponding to the first transcript i.e., half of the dsRNA intensity.

