## Supplemental Tables and Figures for "Capsid Restructuring Activates Semi-Conservative dsRNA Transcription in Cystovirus ɸ6"

#### Supplemental Figure Titles and Legends

**Supplemental Figure 1. Cryogenic electron microscopy of transcribing and transcription-arrested double-layered particles, related to Figure 1.** (**A**) A cryo-EM micrograph of transcribing double-layered particles (tDLP). (**B**) Three representative two-dimensional (2D) class averages of tDLPs show particles with a complete outer P8 layer (top), partially disassembled P8 layer (middle) and fully disassembled P8 layer (bottom). (**C**) A cryo-EM micrograph of transcription-arrested (ta)DLP sample. (**D**) 2D class averages of the particles in the taDLP show similar morphologies to those in the tDLP sample in *B*.

**Supplemental Figure 2. Cryogenic electron microscopy data processing workflow, related to Figures 1–7.** A schematic of the cryo-EM data processing workflow. The point group symmetries applied during reconstruction are given in parenthesis (I, icosahedral symmetry; D3, dihedral with three-fold symmetry; C1, asymmetric). The plots show Fourier shell correlation-based resolution estimation for the transcription-arrested (ta) double-layered (DLP) and single-layered (SLP) particle reconstructions.

**Supplemental Figure 3. P8 shedding from DLP, related to Figure 2.** (**A**) Double layered particles (DLPs) incubated in storage buffer (no NTPs) or transcription buffer with the indicated NTPs for 10 min (the time point for taDLP cryo-EM sample preparation) were analyzed by rate-zonal centrifugation in a 10–30% sucrose gradient and the light-scattering profiles imaged. (**B**) Protein and RNA content of the collected gradient fractions were analyzed by SDS-polyacrylamide and native agarose gel electrophoresis, respectively. The electrophoretic mobility of proteins P1, P2 and P8 (upper and middle panels), and the L, M and S dsRNAs (lower panels) are indicated (left). The middle panels, showing the sedimentation profiles for P8, are also shown in Fig. 2G.

**Supplemental Figure 4. In vitro packaging, replication and transcription activity for ϕ6 aSLP, aSLP$_{\mathrm{lowP2}}$ and aSLP$_{\mathrm{lowP7}}$ related to Figure 3.** (**A**) Protein composition of the *in vitro* assembled SLPs. The *in vitro* assembly reactions for aSLPs were carried out by mixing proteins P1, P2, P4 and P7 in the ratio observed in ϕ6 virions to produce standard aSLPs, or using reduced amount of P2 or P7 in the self-assembly reaction to produce $aSLP_{lowP2}$ and $aSLP_{lowP7}$, respectively. The formed aSLPs were purified using rate zonal centrifugation in a linear 10–30% (wt/vol) sucrose gradient and the collected light-scattering zones were analyzed by SDS-polyacrylamide gel electrophoresis. Purified ϕ6 virions were used as a protein marker. The electrophoretic mobility of relevant ϕ6 proteins is indicated on the left. (**B**) Schematic presentation of the ϕ6 *in vitro* packaging, replication and transcription reaction with unlabeled ssRNA template and $^{33}$P-UTP containing NTP set. The unlabeled genomic (+)ssRNA precursors molecules (blue) are packaged into the empty SLPs (step I) where they are used as templates for minus-strand RNA synthesis catalyzed by the P2 RdRps (step II). The RNA synthesis starts from the 3' terminus of the (+)ssRNA templates and results in the production of $^{33}$P-UTP-labeled negative-strand (orange). After completion of the negative-strand synthesis, the P2 RdRps are positioned in the 3'end of the (-)ssRNA, allowing initiation of the first round of transcription using the newly synthesized negative-strands as template (step III). Due to the semi-conservative nature of ϕ6 transcription, this results in the replacement of the parental unlabeled positive-strands and their release from SLP, while the positive-strands of the dsRNAs become labelled. Thus, both stands of dsRNA are $^{33}$P-labelled at this stage. After the completion of the first round of intra-capsid transcription, the P2 RdRp can re-initiate positive-strand synthesis, resulting in the displacement of the earlier synthesized positive-strand and release of $^{33}$P-labelled (+)ssRNA from the SLP (step IV). Thus, the second and the following (+)RNAs released from SLPs become labeled in this experimental system. After completion of positive-strand synthesis, the RdRp is located at the 5'end of the (-)ssRNA template and requires access to the 3'end of the (-)ssRNA strand to be able to reinitiate transcription (step IV; black arrow). (**C**) Time-course of ϕ6 SLP-directed RNA synthesis. Transcription reactions containing unlabelled ϕ6 s, m and l (+)ssRNAs, $^{33}$P-UTP containing NTP set and the indicated in vitro assembled aSLPs were incubated at 30°C. Aliquots were taken at the indicated time points, analyzed by agarose gel electrophoresis, and the gels were autoradiographed (representative autoradiograms of two to three repetitions are shown). rSLPs produced using recombinant expression system were used as a control. The position of dsRNA (uppercase letters) and ssRNA (lowercase letters) is indicated on the left.

**Supplemental Figure 5. Interaction interfaces between P7 and P1 capsid proteins, related to Figure 5.** Depicted is an inside-out view along the three-fold axis of the transcription arrested single-layered particle (taSLP). (**A–B**) naming of different P7 (**A**) or P1 (**B**) chains. Dashed lines in (**A**) indicate P7 chains not participating in interactions with P1 chains. (**C**) Schematic representation of the curvature-specific elements (CSEs) present on the P7 structure. P7 chains corresponding to different sides of the dimer and the color codes for the three identified CSEs are indicated below the model. (**D**) Different P7—P1 interaction sites are shown on the P1 shell (yellow), with one P7 asymmetric unit removed. The tables show interface residues for each of the interaction sites (a1, a2, b1 and b2) identified by ChimeraX *interfaces* command using area cutoff of 250 Å$^2$. Secondary structure elements of P7s and CSEs (from panel C) are indicated on the P7 surface. The interaction surface area is indicated above each table. Bolded P7 residues indicate interface residues present in the majority (at least 4 of 6) of P7 monomers. Bolded P1 residues indicate common residues between P1 chains involved in the interaction with P7 chains.

**Supplemental Figure 6. P2 RdRp density and model fit, related to Figure 6.** (**A**) Extracted map of P2 RdRp and the corresponding model in stage B, colored by polymerase subdomains. Insets (on the right) show the selected structural elements (β16, β17, α22), the D214-K219 loop that contacts the packaged RNA, as well as the missing loop between α22 and α23 (below), eliminated due to clashes with $P1_A$. (**B**) Outside-in and inside-out views of local resolution estimates of P2 densities in stages A1—A3, B and C with a total enclosed volume of 70,000 $Å^3$ each. Progress through the stages from A1 to C increases the apparent flexibility of P2, especially in the palm and finger subdomains.

**Supplemental Figure 7. The versatility of the P7 α3–α4 basket in interactions with P1 capsid proteins and P2 RdRp, related to Figures 5 and 6.** Maps and models of stage B colored by chain. Central P7 dimer is indicated in green hues in the map, while other P7 dimers are indicated in transparent white. In models, only the central P7 dimer and P1 or P2 chains are indicated. (**A**) Interaction site of P7 with two $P1_B$s. (**B**) Interaction site of P7 at the interface of $P1_A$ and $P1_B$. (**C**) Interaction site of wrapped P7 dimer with P2 and side-interaction with $P1_A$.

### Supplemental Tables and Legends

#### Supplemental Table 1A. Cryo-EM data collection and processing statistics.

| | tDLP-I | tSLP-I | taDLP-I | taSLP-I | taDLP-D3 | taSLP-D3 |
|---|---|---|---|---|---|---|
| Voltage (kV) | 300 | 300 | 300 | 300 | 300 | 300 |
| Movies (no.) | 772 | 772 | 16,177 | 16,177 | 16,177 | 16,177 |
| Electron exposure ($e^{-}/Å^2$) | 50 | 50 | 42 | 42 | 42 | 42 |
| Defocus range (µm) | 1.1–4.0 | 1.1–4.0 | 0.2–4.1 | 0.2–4.1 | 0.2–4.1 | 0.2–4.1 |
| Pixel size (Å) | 2.70 | 2.70 | 1.40 | 1.40 | 1.40 | 1.40 |
| Box size (pixels) | 256 | 256 | 512 | 512 | 512 | 512 |
| Symmetry imposed | I | I | I | I | D3 | D3 |
| Initial particle images (no.) | 5974 | 5974 | 221,143 | 221,143 | 221,143 | 221,143 |
| Final particle images (no.) | 2464 | 2762 | 74,260 | 56,922 | 74,260 | 56,922 |
| Map resolution (Å) | 6.1 | 5.9 | 3.5 | 3.5 | 4.0 | 4.0 |
| FSC threshold | 0.143 | 0.143 | 0.143 | 0.143 | 0.143 | 0.143 |
| Map sharpening *B* factor ($Å^2$) | –437 | –376 | –223 | –211 | –210 | –203 |

#### Supplemental Table 1B. Cryo-EM data collection and processing statistics (continued).

| | taDLP disassembly intermediate 1 | taDLP disassembly intermediate 2 | taDLP disassembly intermediate 3 | taDLP disassembly intermediate 4 |
|---|---|---|---|---|
| Voltage (kV) | 300 | 300 | 300 | 300 |
| Movies (no.) | 16,177 | 16,177 | 16,177 | 16,177 |
| Electron exposure ($e^{-}/Å^2$) | 42 | 42 | 42 | 42 |
| Defocus range (µm) | 0.2–4.1 | 0.2–4.1 | 0.2–4.1 | 0.2–4.1 |
| Pixel size (Å) | 1.40 | 1.40 | 1.40 | 1.40 |
| Box size (pixels) | 512 | 512 | 512 | 512 |
| Symmetry imposed | I | I | I | I |
| Initial particle images (no.) | 221,143 | 221,143 | 221,143 | 221,143 |
| Final particle images (no.) | 11,016 | 6421 | 6909 | 23,578 |
| Map resolution (Å) | 7.1 | 7.9 | 7.5 | 6.3 |
| FSC threshold | 0.143 | 0.143 | 0.143 | 0.143 |
| Map sharpening *B* factor ($Å^2$) | –290 | –312 | –296 | –269 |

#### Supplemental Table 2. Cryo-EM data processing statistic for localized reconstructions of protein layers and P4.

| | DLP inner layer expanded | SLP inner layer expanded | SLP inner layer over-expanded | P4 polar with mRNA from SLP | P4 equatorial from SLP | P4 polar from DLP | P4 equatorial from DLP |
|---|---|---|---|---|---|---|---|
| Parent data set | taDLP-I | taSLP-I | taSLP-I | taSLP-D3 | taSLP-D3 | taDLP-D3 | taDLP-D3 |
| Pixel size (Å) | 1.40 | 1.40 | 1.40 | 1.40 | 1.40 | 1.40 | 1.40 |
| Symmetry imposed | C1 | C1 | C1 | C1 | C1 | C1 | C1 |
| Box size (pixels) | 180 | 144 | 144 | 112 | 112 | 112 | 112 |
| Initial sub-particles (no.) | 4,455,600 | 3,415,298 | 3,415,298 | 341,532 | 341,532 | 445,560 | 445,560 |
| Final sub-particles (no.) | 2,504,986 | 391,662 | 689,037 | 25,297 | 46,586 | 124,691 | 60,535 |
| Map resolution (Å) | 3.5 | 3.7 | 3.5 | 5.9 | 5.2 | 4.8 | 4.8 |
| FSC threshold | 0.143 | 0.143 | 0.143 | 0.143 | 0.143 | 0.143 | 0.143 |
| Map sharpening *B* factor ($Å^2$) | –253 | –219 | –217 | –295 | –243 | –268 | –237 |

**Supplemental Table 3. Cryo-EM data processing statistic for localized reconstructions of capsid internal elements.**

| | **P7 triskelion** | **P2 stage A1** | **P2 stage A2** | **P2 stage A3** | **P2 stage B** | **P2 stage C** |
|---|---|---|---|---|---|---|
| Parent data set | taSLP-D3 | taDLP-D3 | taDLP-D3 | taDLP-D3 | taSLP-D3 | taSLP-D3 |
| Pixel size (Å) | 1.40 | 1.40 | 1.40 | 1.40 | 1.40 | 1.40 |
| Symmetry imposed | C1 | C1 | C1 | C1 | C1 | C1 |
| Box size (pixels) | 232 | 192 | 192 | 192 | 192 | 192 |
| Initial sub-particles (no.) | 257,827 | 445,560 | 445,560 | 445,560 | 445,560 | 445,560 |
| Final sub-particles (no.) | 114,297 | 17,314 | 24,470 | 21,926 | 18,230 | 16,157 |
| Map resolution (Å) | 4.1 | 4.7 | 4.6 | 4.7 | 4.8 | 4.8 |
| FSC threshold | 0.143 | 0.143 | 0.143 | 0.143 | 0.143 | 0.143 |
| Map sharpening *B* factor ($Å^2$) | –180 | –214 | –206 | –203 | –214 | –218 |

**Supplemental Table 4.** Model building and refinement statistics.

| | **P1 shell, expanded with P8** | **P1 shell, expanded** | **P1 shell, overexpanded** | **P1 shell in P2 stage B** | **P7 dimer** |
|---|---|---|---|---|---|
| **Refinement** | | | | | |
| Resolution limit (Å) | 3.5 | 3.7 | 3.5 | 4.4 | 3.8 |
| Model-to-map resolution (Å) | 3.6 | 3.8 | 3.7 | 3.8 | 3.7 |
| FSC threshold | 0.50 | 0.50 | 0.50 | 0.50 | 0.50 |
| Model-to-map CC | | | | | |
| Main chain | 0.77 | 0.79 | 0.81 | 0.66 | 0.76 |
| Side chain | 0.74 | 0.78 | 0.79 | 0.64 | 0.75 |
| Model composition | | | | | |
| Non-hydrogen atoms | 23,399 | 12,219 | 12,219 | 47673 | 1690 |
| Residues | 3051 | 1571 | 1571 | 3064 | 222 |
| *B* factors, mean ($Å^2$) | | | | | |
| Protein | 49.98 | 65.04 | 45.68 | 101.7 | 113.21 |
| **R.m.s. deviations** | | | | | |
| Bond lengths (Å) | 0.008 | 0.008 | 0.007 | 0.006 | 0.008 |
| Bond angles (°) | 0.829 | 1.084 | 1.066 | 0.857 | 1.033 |
| **Validation** | | | | | |
| MolProbity score | 1.94 | 2.06 | 2.04 | 2.22 | 1.67 |
| Clash score | 9.79 | 14.31 | 12.21 | 7.66 | 6.43 |
| Rotamer outliers (%) | 0.16 | 0.15 | 0.23 | 3.62 | 0.00 |
| Ramachandran plot | | | | | |
| Favored (%) | 93.49 | 94.19 | 93.29 | 94.54 | 95.41 |
| Allowed (%) | 6.48 | 5.69 | 6.58 | 5.40 | 4.59 |
| Outliers (%) | 0.03 | 0.13 | 0.13 | 0.07 | 0.00 |

Supplemental Figure 1

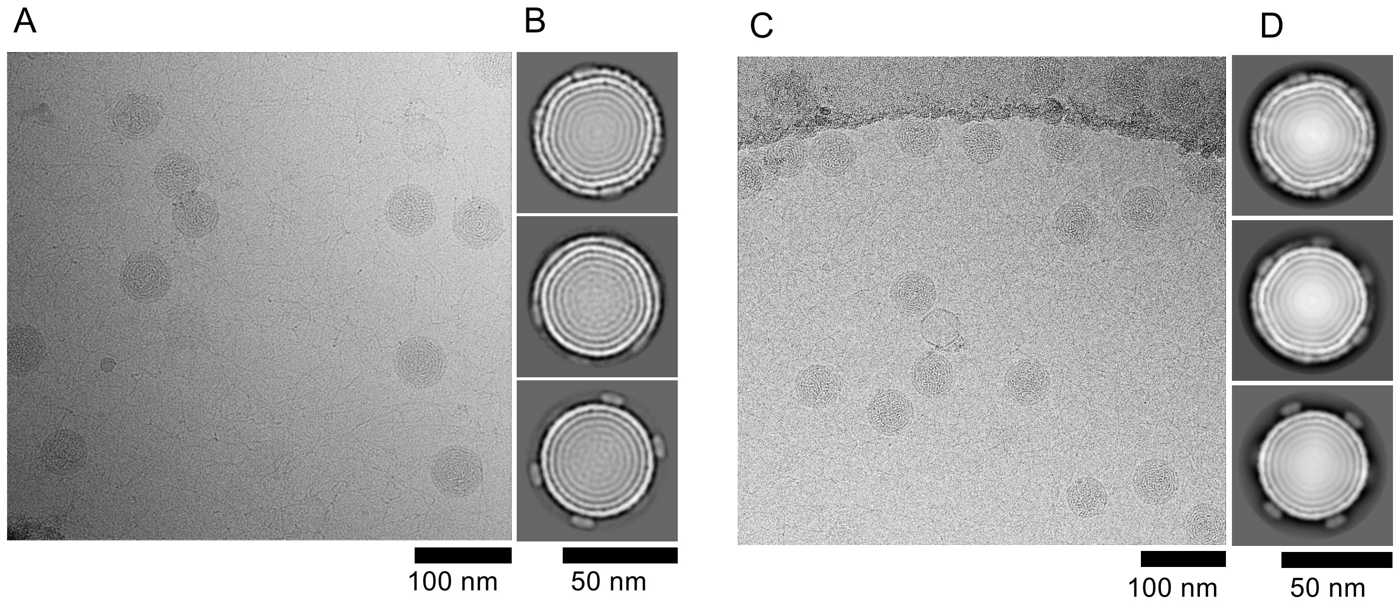

### Supplemental Figure 2

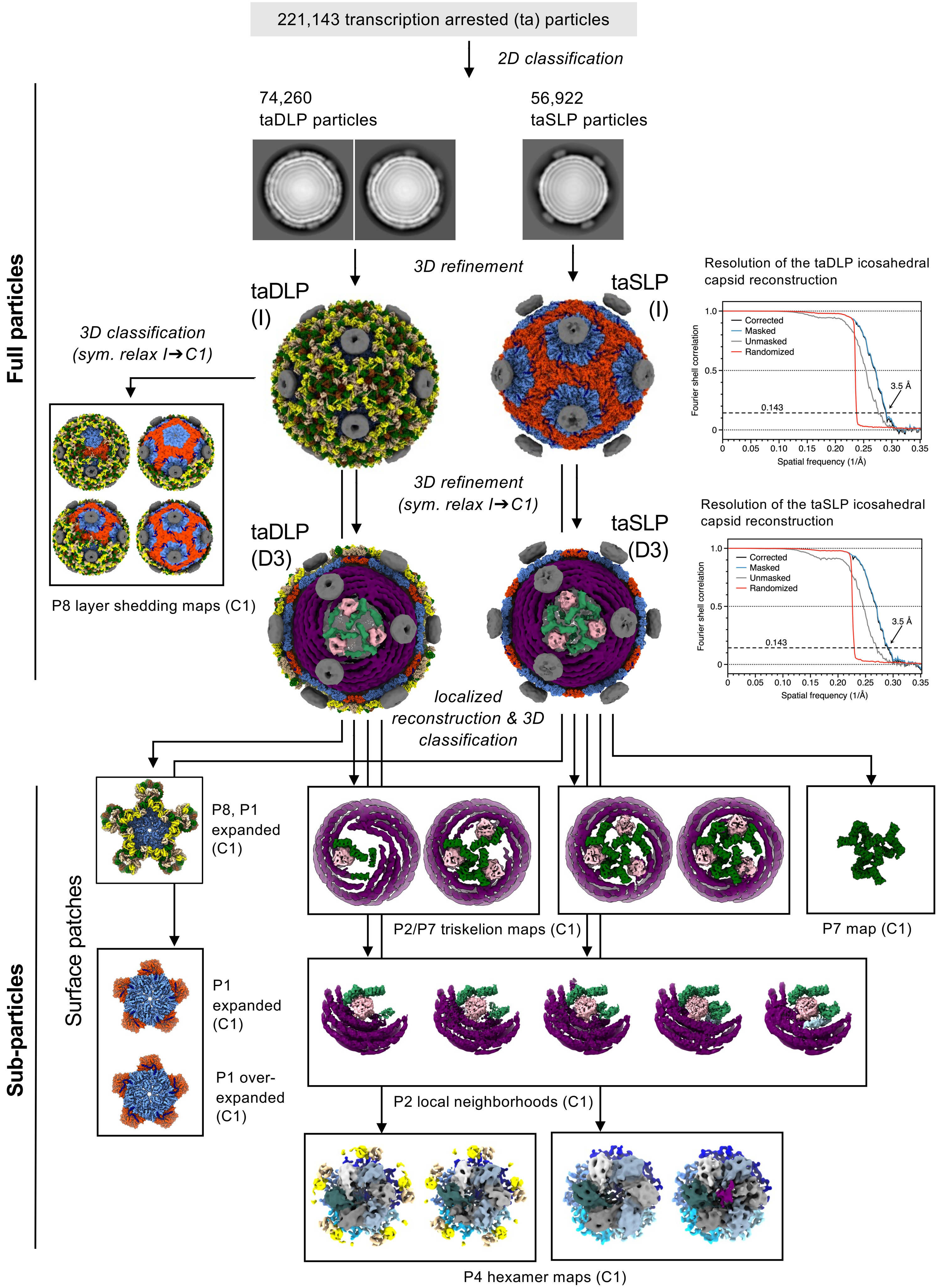

### Supplemental Figure 3

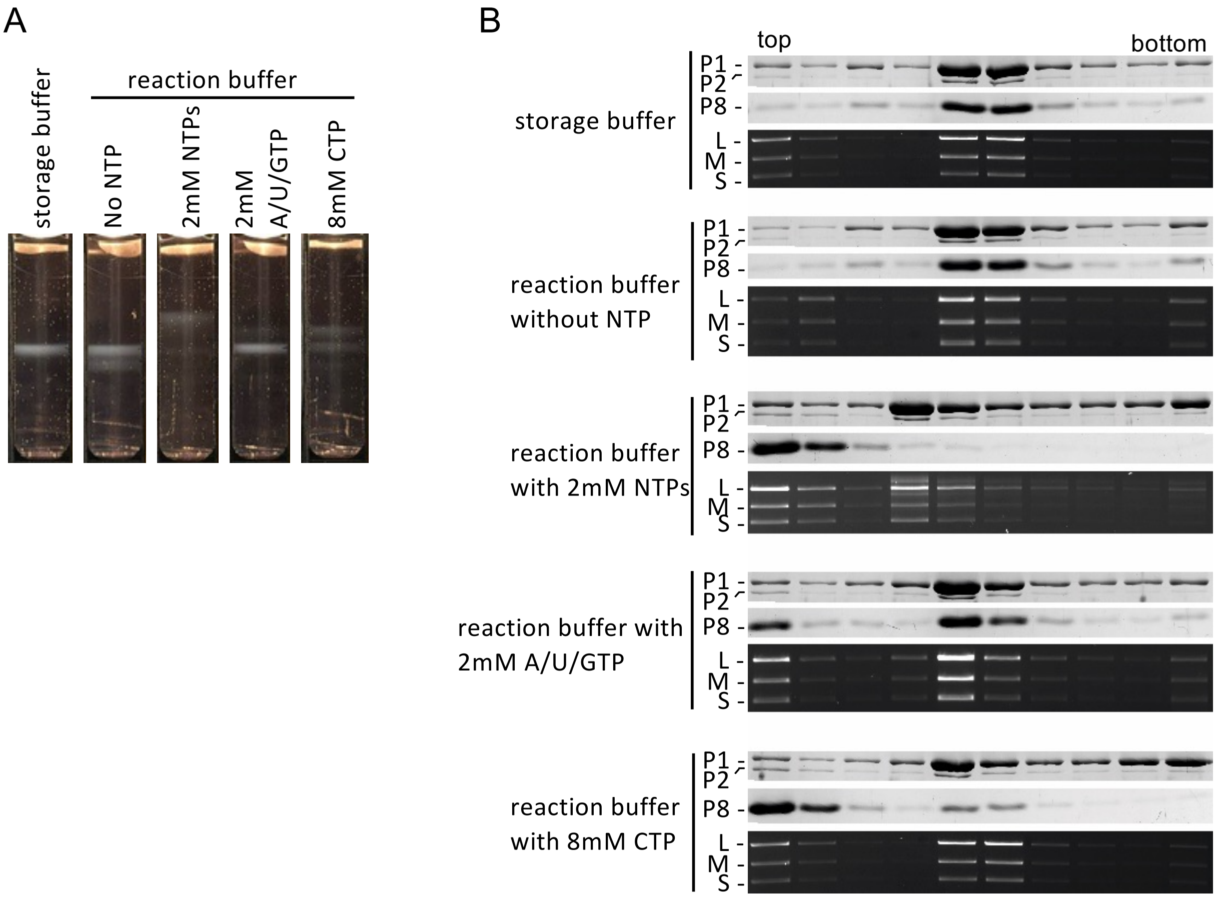

### Supplemental Figure 4

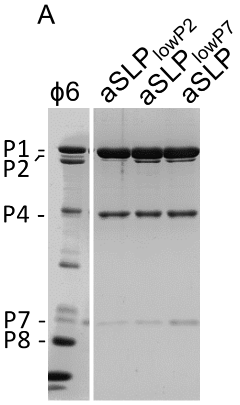


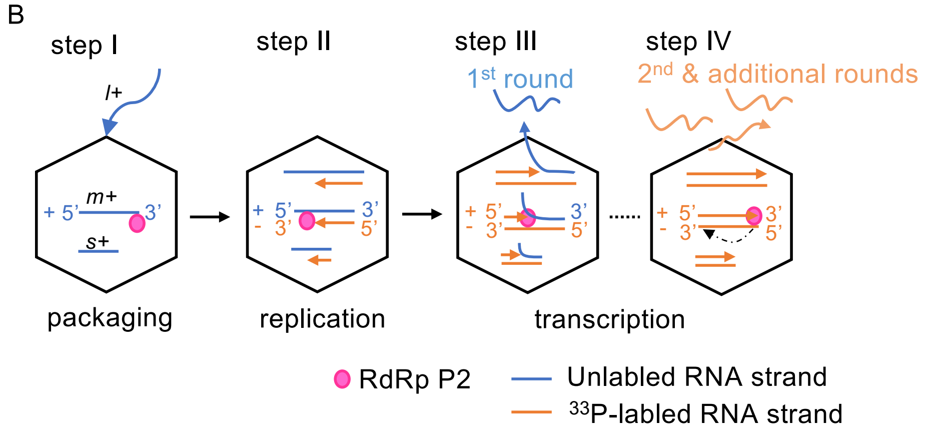


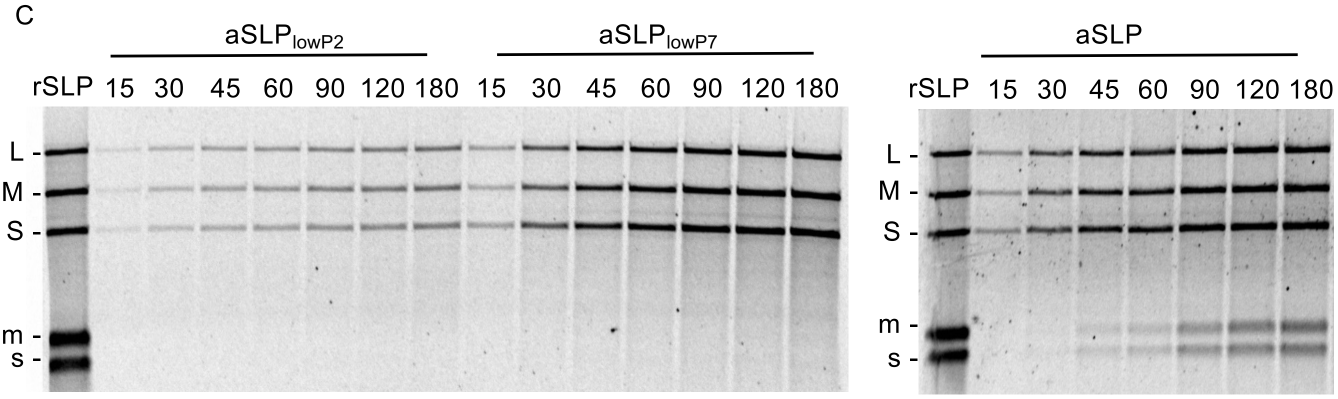

### Supplemental Figure 5

A

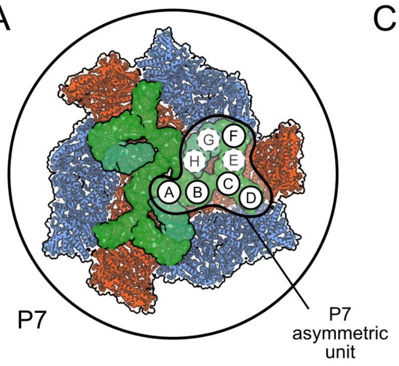


B

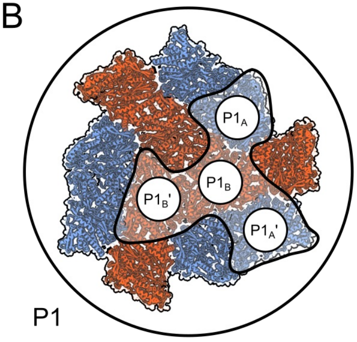


C

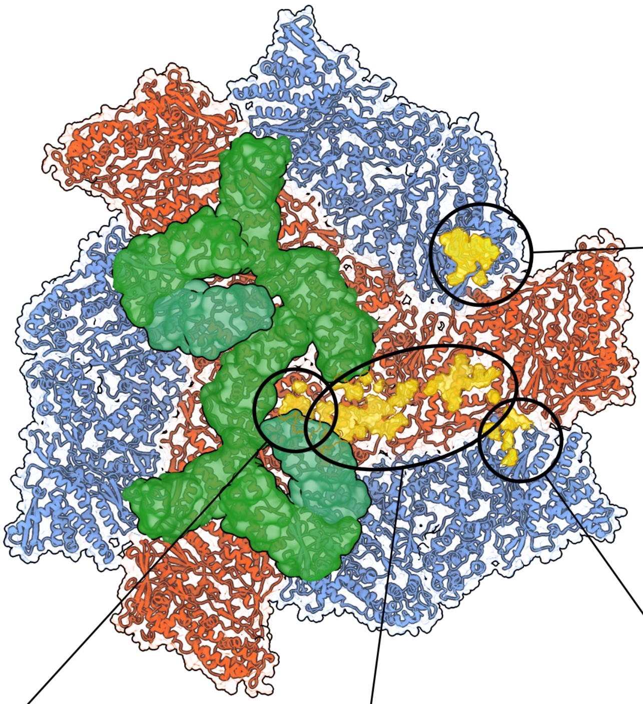

690 Å²

**Site a2**

| P1A | P7F | |
|---|---|---|
| P37 | K14 | CSE1 |
| **Q39** | M74 | |
| **T41** | A75 | CSE2 |
| **R42** | P77 | |
| S272 | Y81 | |
| **N279** | **S83** | α4 |
| **D282** | N86 | |
| Q283 | **H90** | |
| **R285** | D96 | CSE3 |
| L333 | W97 | |
| P335 | I98 | |
| E500 | T99 | |

290 Å²

**Site b2**

| P1B' | P7A | |
|---|---|---|
| **Q39** | D10 | CSE1 |
| **R42** | S11 | |
| **N279** | A75 | CSE2 |
| **D282** | S80 | |
| **Q283** | Y81 | |
| **R285** | R82 | |
| | **S83** | α4 |
| | N86 | |

1700 Å²

**Site b1**

| P1B | | | P7A | | P7B | | P7C | |
|---|---|---|---|---|---|---|---|---|
| T49 | N231 | T683 | N53 | | N59 | | R60 | α3 |
| L52 | T235 | D713 | A56 | | R60 | α3 | H61 | |
| W54 | R239 | R724 | I57 | α3 | H61 | | L69 | |
| E55 | E327 | M758 | N59 | | S63 | | **S83** | |
| N60 | V328 | T762 | R60 | | L69 | | M87 | |
| D62 | T407 | T763 | R82 | CSE2 | **S83** | | I88 | α4 |
| P63 | K564 | I765 | D84 | | N86 | | **H90** | |
| M65 | D567 | | M87 | | M87 | α4 | A91 | |
| H156 | N570 | | I88 | α4 | **H90** | | P93 | CSE3 |
| T200 | H571 | | **H90** | | A91 | | S94 | |
| S203 | Q669 | | A91 | | S94 | CSE3 | | |
| S204 | T673 | | | | | | | |
| K208 | R676 | | | | | | | |
| M209 | M680 | | | | | | | |

310 Å²

**Site a1**

| P1A' | P7D | |
|---|---|---|
| G353 | D10 | CSE1 |
| R523 | S11 | |
| E525 | D13 | |
| R527 | E31 | CSE2 |
| P554 | R82 | |
| Q556 | D84 | α4 |

D

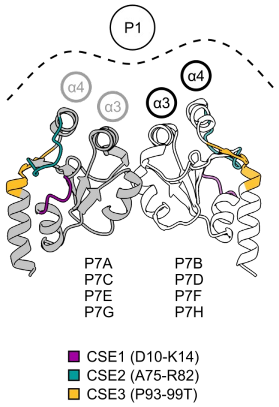

### Supplemental Figure 6

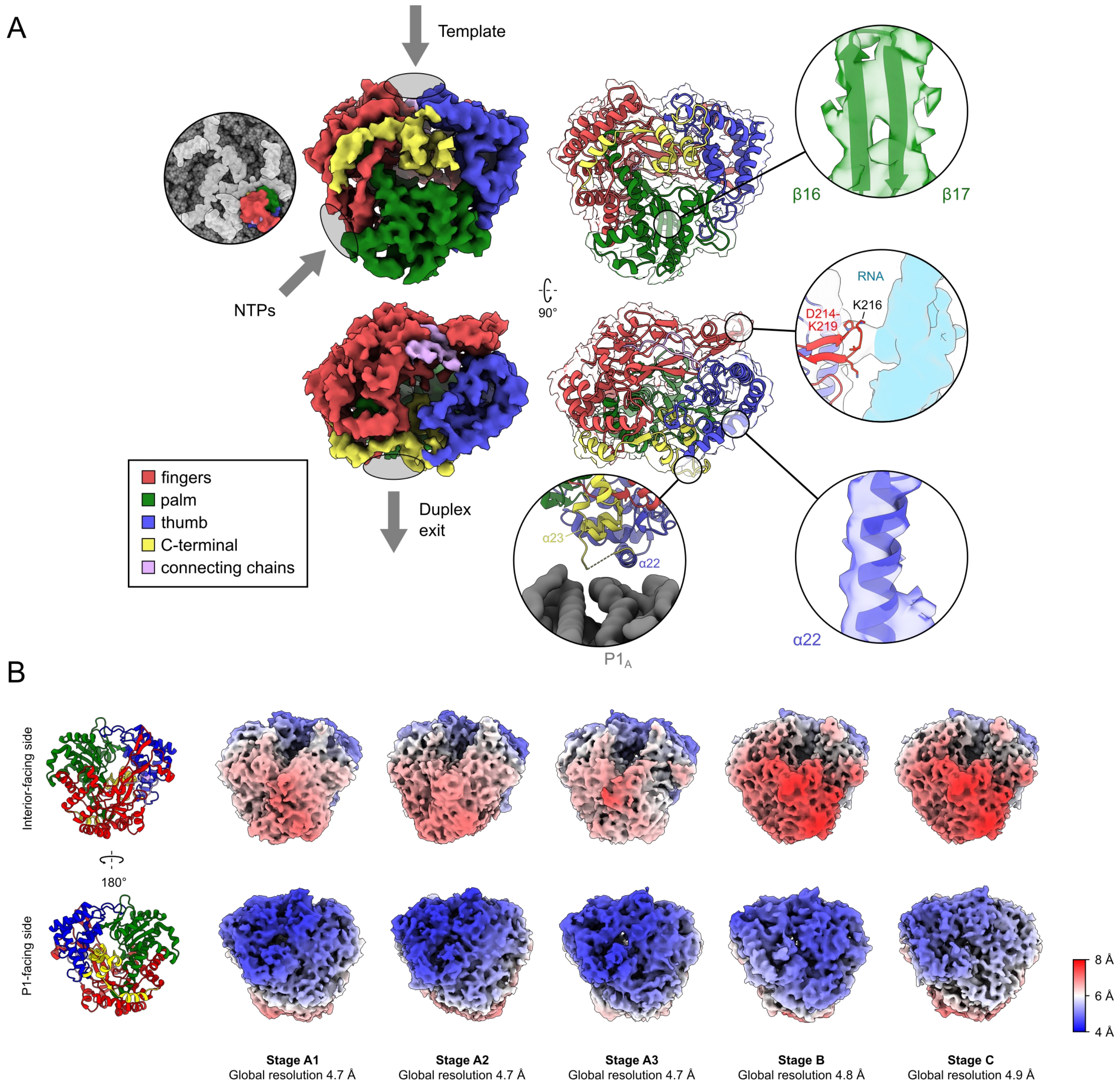

### Supplemental Figure 7

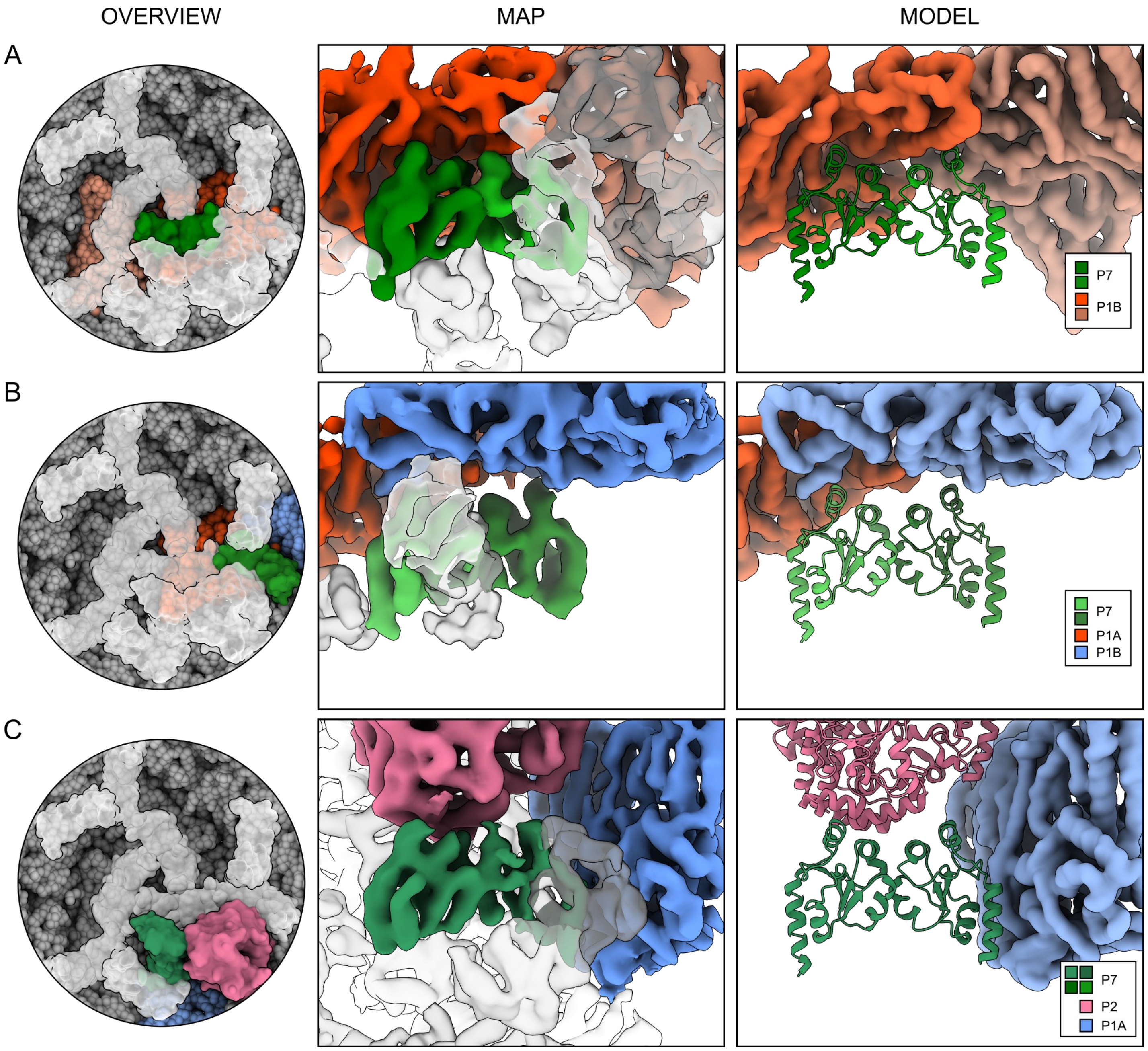
